# Roles for tubulin recruitment and self-organization by TOG domain arrays in Microtubule plus-end tracking and polymerase

**DOI:** 10.1101/340133

**Authors:** Brian Cook, Fred Chang, Ignacio Flor-Parra, Jawdat Al-Bassam

**Author notes:** Correspondence should be addressed to Jawdat Al-Bassam Ignacio Flor-Parra.

## Abstract

The XMAP215/Stu2/Alp14 microtubule polymerases utilize Tumor Overexpressed Gene (TOG) domain arrays to accelerate microtubule plus-end polymerization. Structural studies suggest a microtubule polymerase model in which TOG arrays recruit four αβ-tubulins, forming large square assemblies; an array of TOG1 and TOG2 domains may then unfurl from the square state to polymerize two αβ-tubulins into protofilaments at microtubule ends. Here, we test this model using two biochemically characterized classes of fission yeast Alp14 mutants. Using *in vitro* reconstitution and *in vivo* live cell imaging, we show that αβ-tubulins recruited by TOG1 and TOG2 domains serve non-additive roles in microtubule plus-end tracking and polymerase activities. Alp14 mutants with inactivated square assembly interfaces have defects in processive plus-end tracking and poor microtubule polymerase, indicating a functional role for square assemblies in processive tracking. These studies provide functional insights into how TOG1 and TOG2 domain arrays recruit tubulins and promote polymerase at microtubule plus ends.

## Introduction

Microtubules (MTs) are dynamic polymers that drive diverse cellular functions including mitosis, cytokinesis, cell motility and organelle transport. MTs polymerize from soluble αβ-tubulin dimers (termed αβ-tubulins from herein), via head-to-tail assembly, into protofilaments at MT ends (Akhmanova and Steinmetz, 2015). The polymerization of αβ-tubulins at MT-ends is regulated by GTP hydrolysis, which becomes activated upon the assembly of αβ-tubulins into MT lattices. Activation of GTP hydrolysis at MT ends leads to stochastic dynamic instability transitions where MT-ends switch from polymerization to depolymerization phases (Akhmanova and Steinmetz, 2011, 2015). In cells, conserved classes of MT regulatory proteins regulate different aspects of MT dynamics. One key regulator group is the XMAP215/Stu2/Alp14 family proteins, which accelerate MT polymerization and are universally essential for MT polymerization across eukaryotes. When these MT polymerases are inactivated or depleted, cells generally exhibit fewer MTs, slower MT polymerization rates, and smaller mitotic spindles and chromosome segregation defects. (Akhmanova and Steinmetz, 2015; Al-Bassam and Chang, 2011).

Despite extensive study, a mechanistic understanding for how these MT polymerases function to accelerate MT polymerization or track MT plus-ends is still lacking. XMAP215/Stu2/Alp14 proteins recruit soluble αβ-tubulins via highly conserved N-terminal arrays of Tumor Overexpressed Gene (TOG) domains (Al-Bassam and Chang, 2011) (Brouhard and Rice, 2014). Each array consists of tandem sets of TOG domain classes, TOG1 and TOG2, named according to their position from the N-termini. Yeast MT polymerase proteins, such as Alp14 and Stu2, contain arrays of TOG1 and TOG2 domains followed by the positively charged (SK-rich) domain and dimerize via a C-terminal coiled-coil domain, leading to homodimeric arrays with four TOG domains (Al-Bassam et al., 2006; Haase et al., 2018). The metazoan XMAP215 and human ch-TOG proteins are monomeric with five TOG domain arrays and may form internal dimers with tandem TOG1-TOG2 and TOG3-TOG4 arrays, which are separated by charged regions. These metazoan TOG arrays are comparable to the homodimeric yeast Alp14 or Stu2 TOG1-TOG2 arrays (Al-Bassam and Chang, 2011; Brouhard et al., 2008; Widlund et al., 2011). Recent studies show that TOG domains selectively bind αβ-tubulin in a curved conformation of its soluble form. These findings suggest a potential mechanism for how TOG domains release αβ-tubulins upon adopting a straight conformation during polymerization into the MT plus-ends (Ayaz et al., 2014; Ayaz et al., 2012; Brouhard and Rice, 2014). However, how arrays of TOG domains found in the native MT polymerases function to polymerize αβ-tubulins at the growing MT end still remains unclear.

XMAP215/Stu2/Alp14 MT polymerases localize to the growing plus ends of MTs in a process known as MT plus-end tracking. *In vitro* reconstitution assays demonstrate that these purified proteins are able to track microtubule plus-ends without additional factors (Al-Bassam et al., 2012; Brouhard et al., 2008). Single molecule experiments suggest that single XMAP215 molecules reside at MT plus-ends for many seconds and thus may operate for many rounds of αβ-tubulin polymerization (Brouhard et al., 2008). The mechanisms for MT plus-end tracking by XMAP215/Stu2/Alp14 proteins appear to be distinct from those of other MT plus-end tracking proteins, such as EB1 or CLIP-170 family proteins (Akhmanova and Steinmetz, 2011, 2015; Asbury, 2008; Brouhard et al., 2008). Yeast Alp14 and Metazoan XMAP215 bind tightly to the extreme tips of polymerizing MT plus ends and translocate with them during polymerization (Al-Bassam et al., 2012; Brouhard et al., 2008). In contrast, EB1 proteins bind an extended zone at the MT plus-end defined by the GTP nucleotide state of the polymerized αβ-tubulin, and dissociate without translocation along with polymerizing microtubule plus-ends (Maurer et al., 2011; Maurer et al., 2014). Numerous studies have implicated the TOG domains and charged Sk-rich regions of these proteins in plus end tracking, but the mechanisms for tracking at a structural level remains poorly explored (Al-Bassam et al., 2012; Brouhard et al., 2008; Widlund et al., 2011).

In the accompanying manuscript (Nithianantham et al 2018), we describe structural studies that provide a new mechanistic model for TOG domain arrays and MT polymerase function. We used X-ray crystallography to define two different conformations of TOG1-TOG2 arrays bound to tubulins: 1) a wheel-like square-shaped dimeric assembly in which four TOG domains are bound to four αβ-tubulins in a head to tail fashion, and 2) a putative polymerization complex in which each TOG1-TOG2 subunit is unfurled due to a 68° rotation of TOG2 around TOG1, promoting the concerted polymerization of their bound αβ-tubulins into a curved protofilament. In the polymerized assembly complex, TOG1 and TOG2 specifically bind the lower and upper αβ-tubulins. Our biochemical studies suggest TOG1 and TOG2 exhibit distinct exchange rates for αβ-tubulin. These data suggest a “polarized unfurling” model of MT polymerase function: the square structure may be a recruitment complex that delivers αβ-tubulins in the proper configuration to the protofilament end; there, this complex unfurls its TOG domains promoting the concerted polymerization of two αβ-tubulins onto the growing MT-plus end.

A key question is whether these structures represent true intermediates in MT polymerization pathway. The polarized unfurling model makes specific predictions about the function of distinct TOG domains in recruiting αβ-tubulins and the interfaces between TOG domains, which are responsible for formation of the square assembly. For instance, the model predicts that TOG1 bound tubulin is incorporated at the base of a newly unfurled protofilament to anchor the TOG array, while TOG2 promotes insertion of its αβ-tubulin via rotation around the TOG1 anchored αβ-tubulin. The unfurling process likely requires that TOG2 is anchored onto TOG1 via interface 2. The binding of four αβ-tubulins by the square complex, formed by TOG1-TOG2 dimers and stabilized by interfaces 1 and 2, may serve a critical role in the recruitment, delivery, and pre-arrangement of these αβ-tubulins at the polymerizing MT end (Nithianantham et al 2018).

Here, we study the XMAP215/Stu2/Alp14 MT polymerase mechanism and test this model through analyses of structure-based Alp14 mutants specifically inactivating TOG-tubulin interactions and in the interfaces necessary for square complex assembly (Figure 1-Supplement 1B). We study these mutants using a combination of *in vitro* MT reconstitution assays and *in vivo* imaging of dynamic MTs in S. *pombe* cells. Our studies reveal that αβ-tubulin-recruited by TOG1 and TOG2 domains serve distinct roles in MT polymerase cycle: the TOG1-tubulin interaction is critical for processive MT plus-end tracking, while TOG2-tubulin interaction is critical for accelerating MT polymerization. The assembly of TOG1-TOG2 subunits into a TOG square complex is essential for processive MT-plus end tracking and polymerase activity.

## Results

### The TOG1 and TOG2 recruitment of αβ -tubulin serves distinct roles in MT polymerase and plus-end tracking

TOG domains bind soluble αβ-tubulin via a narrow interfaces formed by the inter-HEAT repeat loops (Nithianantham et al, 2018; Ayaz et al., 2014; Ayaz et al., 2012). Biochemical analyses indicate that the two types of yeast TOG domains exhibit different affinities for αβ-tubulin and these affinities can be modulated by the ionic strength of the solution (Nithanantham et al. 2018; Ayaz et al., 2014). TOG1 binds αβ-tubulin tightly at both 50 mM and 200 mM KCl while TOG2 binds to αβ-tubulin tightly at 50 mM KCl, and loosely at 200 mM KCl (Nithanantham et al. 2018; Figure 1A). To differentiate among the functions of αβ-tubulin recruitment by TOG1 and TOG2 and their contribution to MT-plus end tracking and polymerase activities, we first studied the effect of changing the solution ionic strength from 50 to 200 mM KCl on the MT polymerase activity of dimeric fission yeast Alp14. We used near-native dimeric *S. pombe* Alp14 (residues 1-690; termed wt-Alp14) and dynamic MT polymerization assays visualized by TIRF microscopy (Figure 1 Supplement 2)(Al-Bassam, 2014). Such a change in the ionic strength has virtually no effect on αβ-tubulin polymerization activity at MT plus ends in the absence of Alp14 (Table S1). Our biochemical analyses show that increasing ionic strength from 50 mM to 200 mM KCl rapidly accelerates the αβ-tubulin exchange by the two TOG2 domains in wt-Alp14 (Nithianantham et al, 2018). MT dynamic TIRF assays show that wt-Alp14 accelerates MT plus-end polymerization by roughly four (0.38 μm/min to a maximal 1.52 μm/min) while tightly tracking the tips of MT plus-ends (Figure 1B); (Al-Bassam et al., 2012). In contrast at 50 mM KCl where wt-Alp14 maintains tight αβ-tubulin binding and slow exchange via TOG2 domains, we observe that the maximal MT polymerase for wt-Alp14 plateaus at ∼1.07 μm/min, suggesting a ∼40% decrease in activity compared to its activity at 200 mM KCl (t-test p < 10^−5^).

**Figure 1:**
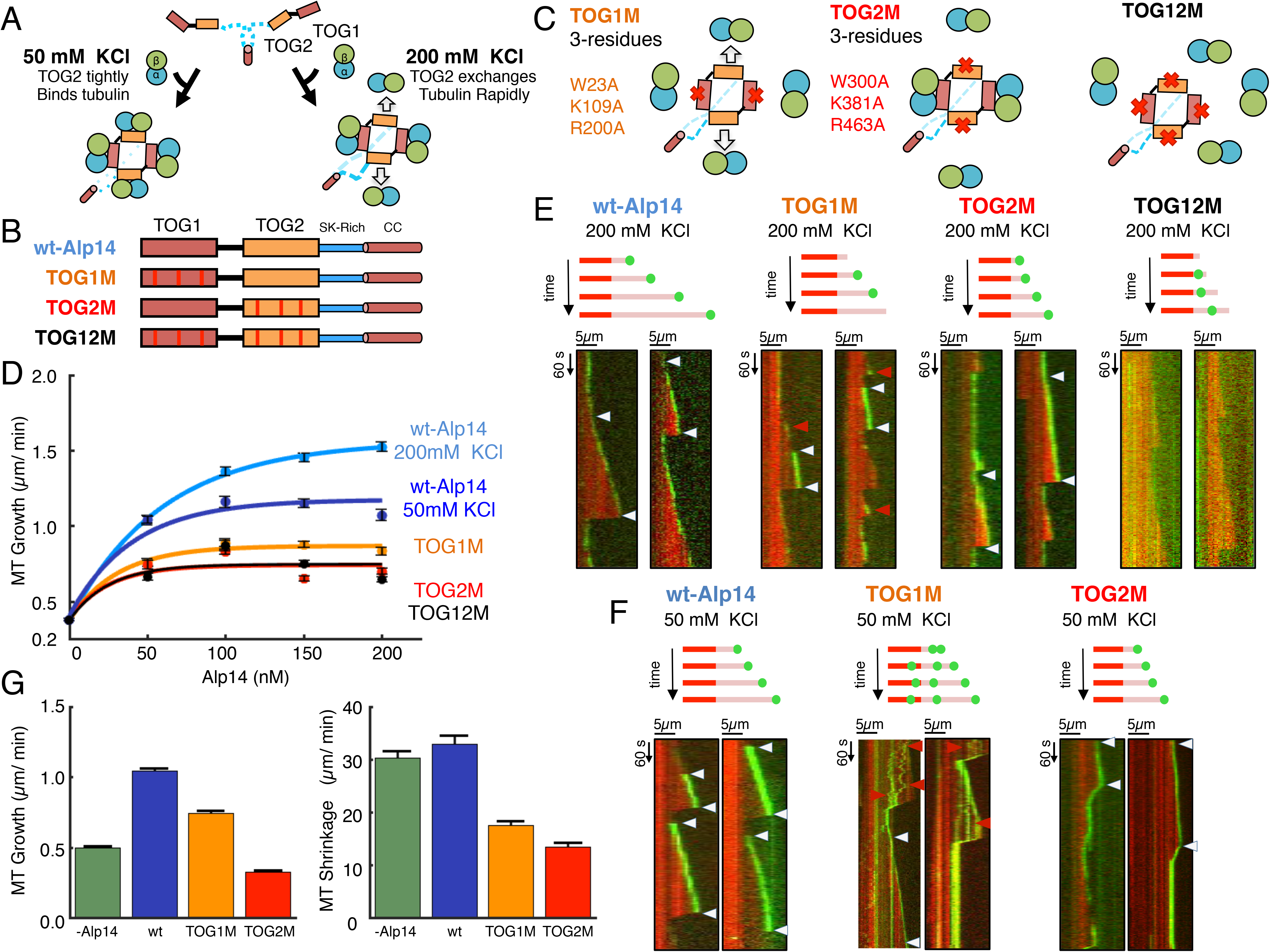
The αβ -tubulin recruited by TOG1 and TOG2 domains serve unique and non-additive roles in MT polymerase and plus-end tracking activities in dimeric Alp14. A) Model for biochemical roles of TOG1 and TOG2 domains in recruiting αβ-tubulins at 50 and 200 mM KCl. TOG1 (red) binds αβ-tubulin (blue and green) tightly at both 50 and 200 mM KCl, while TOG2 (orange) binds αβ-tubulin tightly at 50 mM KCl but exchanges rapidly at 200 mM KCl. B) Near-native Alp14 constructs with αβ-tubulin recruitment defects. Top, wt Alp14, second, TOG1 inactivated (TOG1M); third, TOG2 inactivated (TOG2M), fourth, TOG1-TOG2 inactivated (TOG12M) mutants. C) The activities of TOG1 M, TOG2M, and TOG12M in binding αβ-tubulin as described in (Nithianantham et al, 2018) D) Average MT growth rates for dynamic MTs with various Alp14 constructs. Wt-Alp14 at 200 mM KCl (light blue), wt Alp14 at 50 mM KCl (dark blue). TOG1M (orange), TOG2M (red) and TOG12M (black). The individual values are reported in Table S1. E) Kymographs of individual dynamic MTs with wt-Alp14 or mutants at 200 mM KCl. Top a model for the observed MT polymerization activity and example kymographs shown below. TOG1M shows short MT plus-end tracks (white arrows) or rapid dissociation events (red arrows). TOG2M shows extensive plus end tracking even with non-polymerizing MTs. TOG12M shows no plus-end MT plus end tracking. F) Kymographs of individual dynamic MTs with wt-Alp14 or mutants at 50 mM KCl. Top a model for the observed MT polymerization activity and example kymographs shown below. TOG1M dissociates from MT plus end tracking while dissociating along MT lattices (white arrows). TOG2M binds tightly at MT plus-ends and leads to slow MT depolymerization events. G) Average MT growth and shrinkage rates of dynamic MTs in the absence (-Alp14) and with 50 nM wt-Alp14, TOG1M and TOG2M mutants at 50 mM KCl revealing the uneven contribution of TOG1 and TOG2 to MT polymerase activity.

Next, we studied Alp14 mutants in which αβ-tubulin recruitment by TOG1 and TOG2 is structurally inactivated specifically via mutations in three conserved loops at their αβ-tubulin interfaces (Figure 1C; Figure 1-Supplement 1A-B) (Nithianantham et al, 2018; Ayaz et al., 2014; Ayaz et al., 2012). We studied a TOG1 inactivated Alp14 (termed TOG1M), TOG2 inactivated Alp14 (termed TOG2M) or TOG1 and TOG2 inactivated Alp14 (termed TOG12M) (Al-Bassam et al 2014; Nithianantham et al. 2018). TOG1M contains two active TOG2 domains, while TOG2M contains two active TOG1 domains, and TOG12M does not contain any active TOG domains (Figure 1-Supplement 1A). These mutants were biochemically characterized revealing that TOG2M slowly exchanges αβ-tubulin at both 50 and 200 mM KCl, while TOG1M slowly exchanges αβ-tubulin at 50 mM KCl and rapidly exchanges αβ-tubulin at 200 mM KCl (Al-Bassam et al 2014, Nithianantham et al 2018). We studied the MT polymerase activities of TOG1M, TOG2M and TOG12M at 200 mM KCl using dynamic MT TIRF microscopy assays (Figure 1-Supplement 2A). TOG1M and TOG2M mutants show more than 60-70% decrease in maximal MT polymerase activity, compared to wt-Alp14 (Figure 1D). The maximal MT polymerase activity for TOG1M (∼0.84m/min) is slightly higher than the TOG2M activity (0.70 μm/min) (Figure 1D). Surprisingly, the activity of TOG2M is strikingly indistinguishable from the activity of TOG12M (∼0.65 μm/min)(Figure 1D). Our data suggest the contributions of TOG1 and TOG2 to the Alp14 MT polymerase activity are not additive. Both TOG1 and TOG2 are necessary for different aspects of the MT polymerase mechanism. Although TOG2 exchanges tubulin rapidly, its inactivation leads a more severe loss in MT polymerase activity similar to inactivation of all TOG domains.

To further test whether MT plus-end tracking defects of these mutants relate to differences in the αβ-tubulin exchange rates by TOG1 and TOG2 domains, we studied the MT polymerase activities of TOG1M and TOG2M mutants at 50 mM KCl (Figure 1G). At 50 nM TOG1M, we observe ∼40% decrease in MT polymerase activity (0.74 μm/min) compared to wt-Alp14 (1.07 μm/min), while TOG2M shows a more than ∼83% activity loss (0.38 μm/min), which is lower than the MT polymerization rate in absence of Alp14 (0.50 μm/min). At 50 mM KCl, TOG1M clearly retains higher MT polymerase activity than TOG2M, but a lower activity level than wt-Alp14. At 50 mM KCl, MT shrinkage was dramatically decreased for TOG1M and TOG2M in contrast to wt-Alp14, which showed a similar rate to the absence of Alp14 (Figure 1G).

Next, we analyzed the MT plus-end tracking activities for wt-Alp14 and the three mutants at 200 mM KCl (Figure 1E). wt-Alp14 tightly tracks polymerizing MT-plus ends throughout MT polymerizing phases. In the presence of 200 nM wt-Alp14, almost all MT ends retain plus-end tracking signal throughout polymerization phases (as seen in Figure 1-Supplement 2B) (Al-Bassam et al., 2012). At saturating 200 nM conditions, Alp14 exhibits long tracks at polymerizing MT plus-ends (Figure 1E, left panel). In comparison, TOG1M only partially tracks polymerizing MT plus-ends and shows mostly short tracks, with extensive periods without TOG1M tracking at MT plus ends (Figure 1E, second panel). In contrast, TOG2M tracks polymerizing MT plus-end in a similar manner to wt-Alp14, with long tracks at MT plus-ends throughout polymerizing phases (Figure 1E, third panel). TOG12M does not bind polymerizing MT-plus ends, and mostly diffuses along MT lattices through polymerizing phases and was thus not studied any further *in vitro* (Figure 1E, fourth panel).

At 50 mM KCl, TOG2M and TOG1M mutants track MT-plus-ends with unique patterns indicative with differences in TOG domain affinities to MT plus-ends. Both wt-Alp14 and TOG2M tightly track the tips of polymerizing MT plus-ends. In contrast, TOG1M localization at growing MT plus-ends changes dramatically from 200 to 50 mM KCl (Figure 1F-left and third). TOG1M molecules regularly dissociate from MT plus-ends and diffuse along lattices during MT growth phases, and molecules may rebind MT-plus ends during MT shrinkage phases (Figure 1F-second panel). TOG2M, in contrast, remains tightly bound to growing and shrinking MT plus-ends (Figure 1F-third panel). These data support the idea that the αβ-tubulins recruited by TOG1 and TOG2 serves distinct roles in MT-plus end tracking and MT polymerase activities. Thus, TOG1 is more essential for MT plus-end tracking, while TOG2 serves a greater role in MT polymerase activity. The roles of TOG1 and TOG2 domains within Alp14 arrays are not equal or additive for these functions. Our studies suggest that the αβ-tubulin recruitment by TOG1 and TOG2 domains play unequal roles in MT plus-end tracking and polymerase activities of Alp14.

### The αβ -tubulins recruited by TOG1, but not TOG2, domains are critical for processive Alp14 MT plus-end tracking

To understand the role of TOG1 and TOG2 αβ-tubulin recruitment during MT plus-end tracking, we measured the average residence ratios and their variation for wt-Alp14, TOG1M and TOG2M at MT plus-ends in saturating conditions (200 nM Alp14) using dynamic TIRF assays. For each tracking event, the residence ratio is determined as the duration of Alp14 plus-end signal tracks compared to duration of MT-plus end polymerizing phase (Figure 2A, further detail in Figure 2 Supplement 1; see materials and methods). Figure 2B shows that wt-Alp14 associates with polymerizing MT-plus ends throughout polymerizing phases and exhibits a very high residence ratio and low variance (89.42% average; range 36.9%). In contrast, TOG1M binds MT-plus ends with a lower residence ratio and high variability (72.37% average; range 86.13%) (Figure 2B). TOG2M associates with MT plus-ends with a high residence ratio and low variance, nearly identical to wt-Alp14 (average 91.1%; range 40%) (Figure 2B). The degree of variance in MT plus-end tracking ratios at 200 nM Alp14 suggests that multiple Alp14 molecules may be synchronized at MT-plus ends or form cooperative interactions at MT plus ends and in the TOG1M these events become asynchronous. Thus, TOG1 is essential for MT plus-end association leading to long plus-end tracks, while TOG2 domain inactivation does not affect tracking.

**Figure 2:**
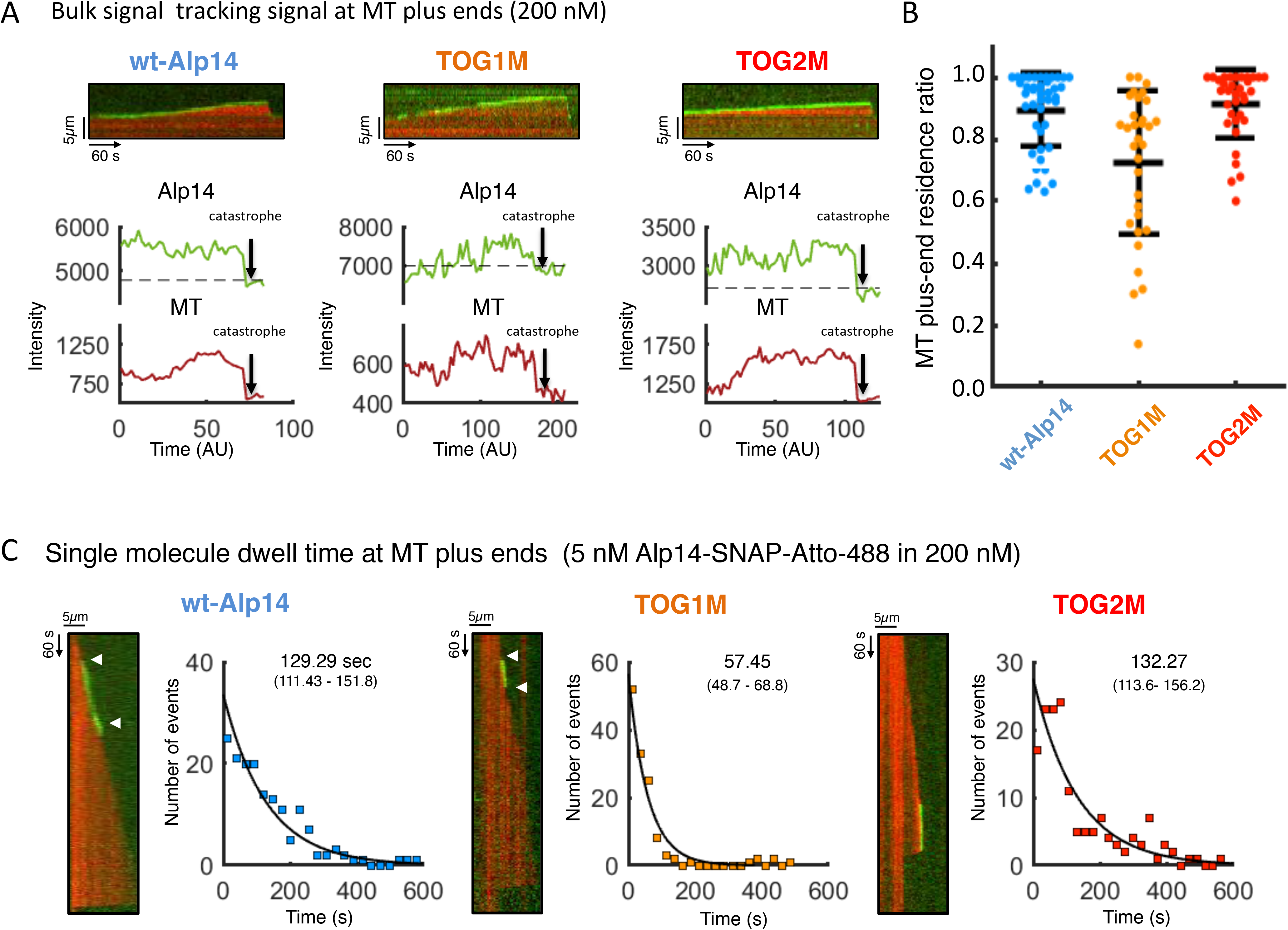
TOG1 and not TOG2 domains are critical for processive MT plus-end tracking. A) Examples of measuring tracking ratio for wt Alp14, TOG1M, TOG2M at growing MT plus ends. Top panels, horizontal kymograph image, Middle panel, Alp14 plus end tracking intensity (green) compared to background (broken line). Lower panel, intensity of MT intensity (red) at the plus end during the polymerizing phase till catastrophe (arrow) compared to background (broken line). wt-Alp14 and TOG2M bulk signal is found at MT plus-ends throughout the process, while TOG1M shows short zones of no signal denoted by decrease in intensity, below the background levels. Additional examples are found in Figure 2 Supplement 1. B) Averages and distributions of tracking ratios for wt Alp14, TOG1M and TOG2M showing a tight distribution for wt-Alp14 and TOG2M, while TOG1M shows a high variability with lower average tracking suggesting poor plus end association. C) Single molecule dynamic MT polymerization studies reveal the average dwell time for wt-Alp14 (blue), TOG1M (orange) and TOG2M (red) mutants. Each panel consists of Left, example kymographs, dwell time frequencies, fit to an exponential decay model. Note that wt-Alp14 is processive while TOG1M has one-third the dwell time, and TOG2M has a similar association to wt Alp14. Details of these single molecule experiments and additional single molecule events are found in Figure 2 Supplement 2 .

Next, we utilized single molecule MT dynamic TIRF assays to determine the average dwell time of Alp14 molecules at MT-plus ends. We used C-terminally SNAPf-tagged Alp14 (termed Alp14-SNAP from herein) on which SNAP-surface Atto-488 fluorophores (see Materials and Methods) is covalently attached, without affecting their MT regulatory functions (Figure 2-Supplement 2A see materials and methods). To visualize the binding and dissociation of single molecules, we diluted 5-10 nM Atto-488 labeled Alp14-SNAP into 190-195 nM non-labeled Alp14-SNAP (Figure 2-Supplement 2A). In these conditions, very few fluorescent Alp14-SNAP-Atto-488 molecules bind to polymerizing MT-plus ends (Figure 2-Supplement 2). We confirmed these are single fluorophores through intensity shifts and photobleaching profiles (Figure 2-Supplement 2C-D). To measure dwell times of individual fluorophores, 100-150 events were fitted to an exponential decay model to determine the average dwell time (Figure 2C; see Materials and Methods). We measured a dwell time of 129.3 sec (n=164) for wt-Alp14 at polymerizing MT plus-ends, suggesting it is an extremely processive tracking protein (Figure 2C; additional example kymographs are shown in Figure 2-Supplement 2C). Alp14 molecules are 20-fold more processive at tracking MT plus ends than XMAP215 molecules. This difference is likely due the presence of two TOG1 and TOG2 domains in each Alp14 molecule compared to only active TOG1 and TOG2 domains per XMAP215 molecule (Brouhard et al., 2008). We measured an average dwell time for TOG1M molecules of 57.4 sec (n=130), which is roughly one-third that observed for wt-Alp14 (Figure 2C; additional example kymographs are shown in Figure 2-Supplement 2D). TOG2M shows an average dwell time of 132.2 sec (n=152) at MT plus ends similar to wt-Alp14 (Figure 2C; additional example kymographs are shown in Figure 2-Supplement 2E). These studies show that at a single molecule level, TOG1M displays short MT plus-end dwell time, which is consistent with the defects observed in MT plus end residence ratios in bulk MT dynamic assays (Figure 2A-B). Our studies suggest that TOG1 domain is critical for processive MT plus-end association, while TOG2 domain is critical for MT polymerase activity, but its inactivation leads to no defect in MT plus-end tracking, in the presence of TOG1 domains. Using the single molecule dwell times and MT polymerase rates, we calculated the tubulin subunits added per MT-end or protofilament in each binding event, as described in Table S2. The wt-Alp14 polymerizes ∼422 αβ-tubulin subunits per event, TOG1M promotes ∼99 αβ-tubulin subunits per event, while TOG2M promotes the polymerization of ∼193 αβ-tubulin subunits due to its longer dwell time.

### Inactivating αβ-tubulin recruitment by TOG1 and TOG2 domains *in vivo* reveals distinct defects in Alp14 MT polymerase and plus-end tracking activities

We next studied the activity of these mutants in living fission yeast cells. The effects of Alp14 on MT dynamics in interphase and mitosis have been extensively characterized in fission yeast (Al-Bassam et al., 2012; Kakui et al., 2013; Okada et al., 2014; Flor-Parra et al., 2018). We replaced the chromosomal *alp14*+ gene with TOG1M, TOG2M, or TOG12M Alp14 mutants and expressed them from the endogenous *alp14* promoter. A similar set of C-terminally GFP-tagged Alp14 genes were also generated (Al-Bassam et al., 2012)(Figure 3-Supplement 1A).

First, we determined the effects on these mutants on Alp14 function on yeast colony growth using spot assays. *alp14*Δ cells are known to be sensitive to growth in cold (20 °C) and high temperatures (36 °C) (Garcia et al., 2001) and to the MT inhibitor Methyl 2-benzimidazolecarbamate (MBC)(Sato and Toda, 2007; Sato et al., 2004). TOG12M cells showed similar growth defects as the null strain (Figure 3-Supplement 1A). Surprisingly, TOG1M cells showed little defect with or without MBC at 20-36 °C (Figure 3-Supplement 1A). In contrast, TOG2M cells displayed a more severe growth phenotype compared to TOG1M cells, which is intermediate between wt and TOG12M cell phenotypes at 20-36 °C (Figure 3-Supplement 1A). These results show that TOG2M and TOG12M exhibit defects at the level of cell proliferation.

Next, we determined the abilities of these mutants on MT polymerase and plus-end tracking activities on interphase MTs. We imaged yeast strains expressing Alp14-GFP and mCherry α-tubulin (example fields shown in Figure 3-Supplement 1B) and analyzed time lapse sequences and corresponding kymographs (Figure 3A, for details on approach see Figure 3-Supplement 2). MT polymerization rates in *alp14* null cells (1.1 μm/min) was about 2-3 fold lower than in *alp14*+ cells (3.18 ± 0.85 μm/min), similar to as described previously (Al-Bassam et al., 2012; Flor-Parra 2018)(Table S3; Figure 3A-B). TOG1M and TOG2M showed significant decreases in MT polymerization rate (1.55 ± 0.48 μm/min and 1.32 ± 0.31 μm/min, respectively), while the TOG12M strain showed a more severe growth defect similar to *alp14*-null cells (1.15 ± 0.25 μm/min). These results contrast with a previous result that a deletion of the TOG1 domain only gives a small defect in MT growth rate (Flor-Parra et al., 2018); it is not clear why this deletion mutant of TOG1 has less of an effect than the TOG1M point mutant, but one reason could be that the TOG1Δ protein assembles into an abnormal complex that nevertheless is functional. *alp14* null mutants also affect MT shrinkage rates (Al-Bassam et al., 2012), and consistently, the TOG mutants affected shrinkage to a similar degree as the polymerization rate (Figure 3-Supplement 3A). TOG12M shows a three-fold decrease compared to wt cells, while TOG1M and TOG2M show 30 and 90% MT shrinkage rate decrease, respectively, with TOG2M being more severe than TOG1M and similar to TOG12M (Figure 3-Supplement 3A)(Al-Bassam et al., 2012). Thus, these results show that these TOG mutations cause significant defects in MT polymerase activity.

**Figure 3:**
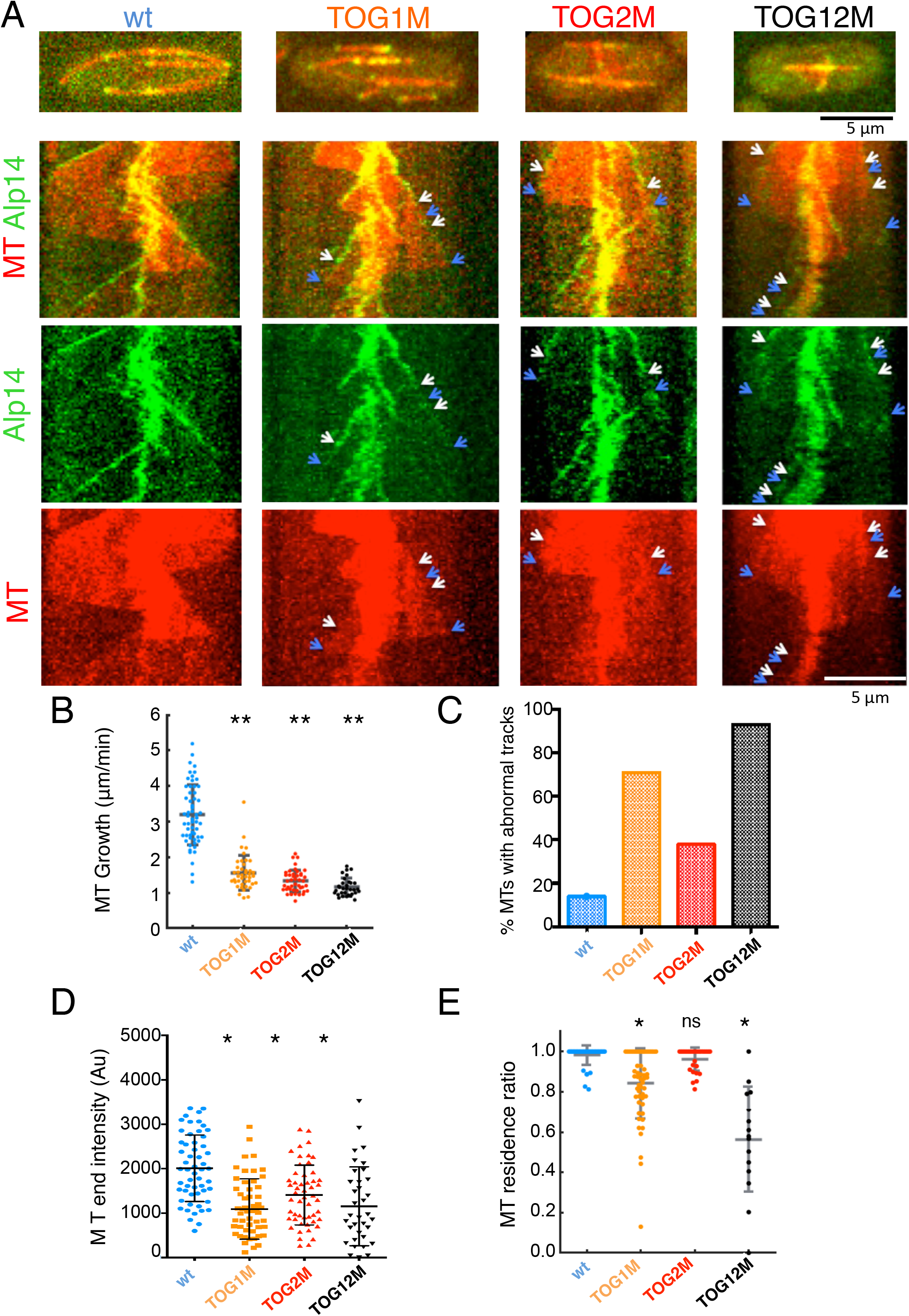
TOG1 and TOG2 domains serve unique functions in MT polymerase and plus-end tracking *in vivo*. A) Images of S. *pombe* cells expressing wt-Alp14, TOG1M, TOG2M and TOG12M mutants. Top, raw cell images with MTs marked with mCherry-Atb2 tubulin (red) and Alp14-GFP (Green). Below, Kymographs of dynamic MTs and Alp14-GFP and the isolated Alp14-GFP data. Note wt-Alp14 tracks MT plus end. TOG1 M, TOG2M and TOG12M all track the plus ends of dynamic MTs. Note the changes in dynamic MT polymerization rates as shown from slopes of lines in the Alp14 kymographs. White arrows mark the beginning of abnormal MT polymerization event, and blue arrow mark the end of such events. Additional example kymographs are shown in Figure 3-Supplement 3D B) MT growth rates of wt-Alp14, TOG1M, TOG2M and TOG12M showing function of TOG domains in MT polymerization (Table S3). Note TOG2M is similar to TOG12M while TOG1 M is shows slightly higher MT growth rate. MT lengths and shrinkage rates are shown in Figure 3-Supplement 3B. C) The proportion of dynamic MTs with abnormal tracking behavior in each strain. Note TOG1 M, TOG2M and TOG12M have higher ratio of these events compared to wt cells. D) The distribution and average of Alp14-GFP intensities at growing MT plus-ends in each of the strains. Note that all the other strains show lower intensity at MT plus-ends. E) MT plus-end Residence / persistence tracking ratios of Alp14 signal at dynamic MTs. The wt-Alp14, TOG2M show high tracking ratio (80-100%) while TOG1 shows a lower and highly variable MT-plus end tracking ratio (10-70%) and TOG12M shows a very low plus end tracking ratio (10-60%). The average values are reported in Table S1. The ** indicates differences that are significant to values of p <10^−5^.

The TOG1M and TOG12M mutants also exhibited defects in MT plus end tracking (Figure 3A, see additional example kymographs in Figure 3-Supplement 3D). In general, the TOG1M, TOG2M and TOG12M mutant Alp14 proteins still localized to growing MT plus ends to varying degrees. Measurements of fluorescence intensity of Alp14 and mutants tagged with GFP showed that TOG1M, TOG2M and TOG12M exhibited significant decreases in intensities with increased variability at MT plus ends (Figure 3D; Table S3). Visual examination of movies and kymographs revealed examples of loss of persistent end tracking (see arrows in Figure 3A; see additional example kymographs in Figure 3-Supplement 3D). In contrast to wildtype cells in which Alp14-GFP intensity was maintained in the large majority of MTs, we found many MTs in TOG1M and TOG12M mutants in which Alp14-GFP fluorescence intensity abruptly decreased or was lost all together at the end of a growing MT (Figure 3D). In contrast there was no change in Alp14 signal in TOG2M MT plus-ends, albeit the signal is generally lower than wt cells. (Figure 3A, see additional examples in Figure 3 Supplement 3D).-Periods in which Alp14 tracking signal came on and then off the plus end were also occasionally observed (Figure 3A; Figure 3-Supplement 1D). Visual inspection of kymographs showed that roughly 71% of MTs in TOG1 (n=55) and 90% in TOG12 mutants (n=14) exhibited some abnormality in maintenance of Alp14 intensity (Figure 3E, Figure 3 Supplement 2, 3D).

Similar to the *in vitro* analyses, we measured the Alp14 residence ratio for individual MT polymerization events, counting time periods in which GFP fluorescence intensity was visually absent or much dimmer while the MT continued to grow. Plots showed normal tracking ratios in WT and TOG2M mutants (Figure 3E; wt: average 98.1 %; range 18.7%; TOG2M: average 96.1%; range 18.7%; no statistically significant difference), extensive variability in TOG1M (Figure 3E; average 84.2%; range 87.0%) and further decreased tracking ratio and increased variability in TOG12M mutant (Figure 3E: average 56.43%; range 100%). Collectively, these measurements revealed that TOG2M has largely normal tip tracking with significant decrease in tracking intensity, while TOG1M and TOG12M mutants have significant tracking defects and decrease in intensity, with the TOG12M mutants showing some tracking defect on almost every MT measured (Figure 3C-E). We also studied the MT bundle length distribution *in vivo* (Figure 3-Supplement 3A) and the proportion of MTs reaching the cortex prior to undergoing catastrophe (Figure 3-Supplement 3C) and observed defects in all the mutants suggesting shorter average dynamic MT lengths and fewer MTs reaching the cell cortex probably due to poor polymerization rates leading to shorter lengths. In sum, these data are consistent with defects seen *in vitro* and show that both TOG domains are needed for optimal MT polymerase and tracking activity *in vivo*. Further, TOG2M has a more critical role in MT polymerase activity, while TOG1 is more critical for MT plus end tracking.

### TOG square assembly interfaces are critical for MT polymerase activity

We present X-ray structures in the accompanying manuscript suggesting that TOG1-TOG2 arrays self-assemble into dimeric square assemblies via two sets of interfaces 1 and 2 (Nithianantham et al, 2018). To test the function of the square assembly conformation, we generated and studied three Alp14 mutants that specifically disrupt the interfaces that hold together the square assembly organization (Figure 4A-B, Figure 1-Supplement 1A-B). We generated an 8-residue mutant that inactivates interface 1, which is responsible for stabilizing the inter-subunit dimerization interface (termed INT1). We generated a 7-residue mutant that inactivates interface 2, which stabilizes the intra-subunit interface (termed INT2). We also constructed a dual interface mutant that mutates all 15-residues to inactivate both intra- and inter-subunit interfaces (termed INT12) (Figure 4A-B). Biochemical analyses show that these mutants behave nearly identical to wt-Alp14 in terms of tubulin binding: they bind αβ-tubulin with near identical stoichiometry, and exhibit rapid exchange by their TOG2 domains (Nithianantham et al, 2018). As predicted, these interface mutants display a conformational destabilization of their square organization. Negative staining electron microscopy reveals wt-Alp14 complexes form compact organizations while INT mutant complexes form abnormal extended organization. The INT2, and INT12 mutants formed either disorganized “necklace-like” array composed of 8-nm densities, while INT1 formed mostly 16-nm filament like extended assemblies. Thus, INT12 and INT2 likely consist of flexibly arrays of TOG domains bound to multiple αβ-tubulin, each of which is 8-nm, while INT1 mostly forms spontaneously polymerized array composed of two polymerized αβ-tubulin assemblies, each of which is 16-nm in length (Nithianantham et al 2018).

**Figure 4:**
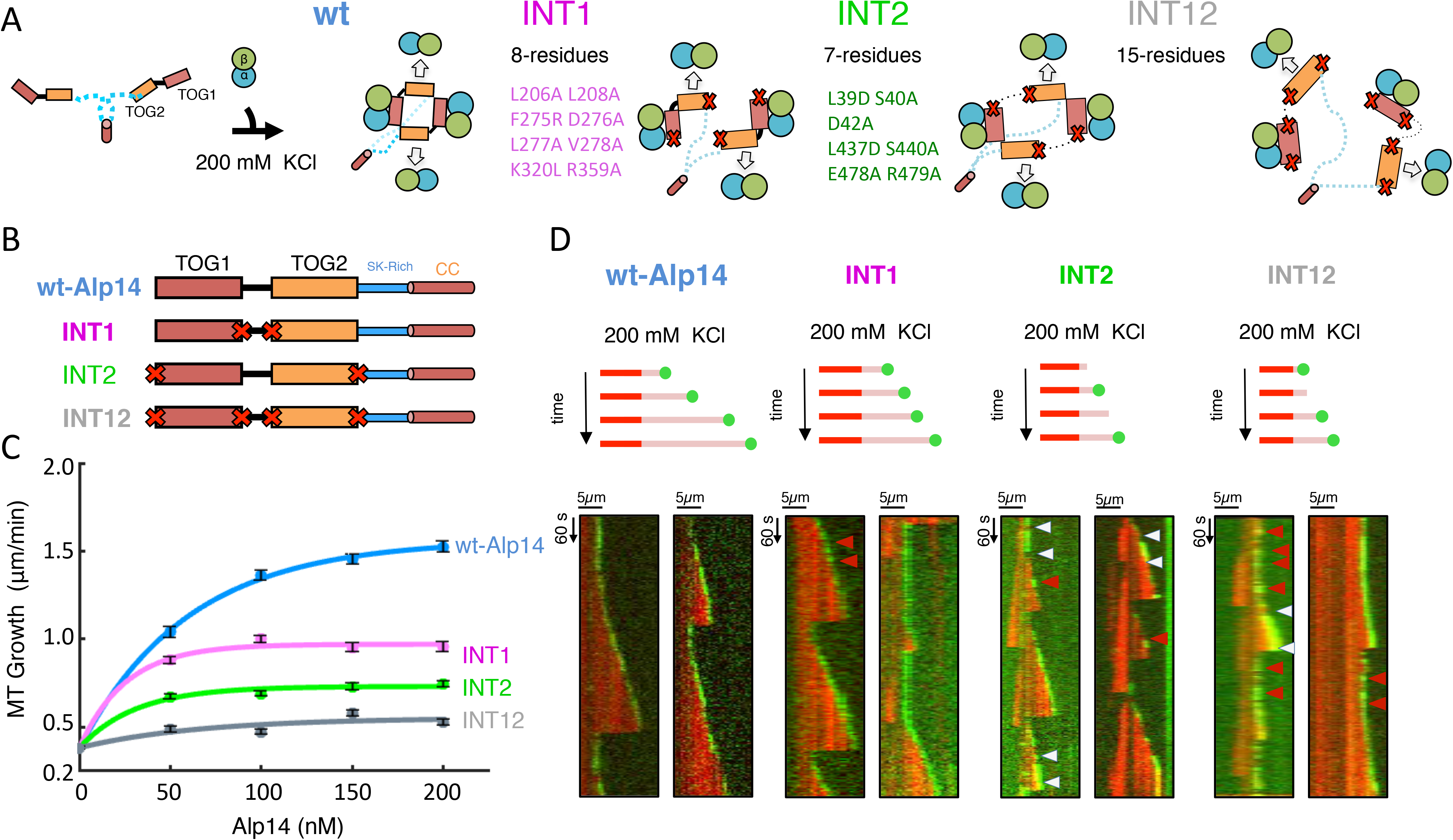
TOG square assembly interfaces are critical for MT polymerase and plus end tracking activities. A) Model for biochemical activities of TOG square assembly interface mutants (as described in Figure 1-Supplement 1A-B and explored in Nithianantham et al 2018). Left, wt-Alp14 forms a square assembly with TOG1 binding tightly to tubulin and TOG2 exchanging tubulin rapidly. INT1 mutant has a defect in dimer assembly through destabilizing interface 1. INT2 mutant has a defect in TOG2 interface with TOG1 in each subunit while still maintaining interface 1. INT12 has defects in both interfaces 1 and 2 leading to destabilized TOG array without defects in αβ-tubulin binding. B) Alp14 constructs with defects TOG square assembly interfaces. Alp14 includes TOG1 and TOG2 domains, SR-rich region (SR-rich) and coiled-coil (cc) domain. Top, wt Alp14, second, INT1 has inactivated interface 1 through introducing eight-residue mutations Third, INT2 inactivated interface2 through seven-residue mutations. Fourth INT12 includes inactivation of both interfaces 1 and 2. C) MT polymerase activities of wt, INT1, INT2, INT12 mutants at 200 mM KCl. INT1 shows a 60% decrease in MT polymerase, while INT2 and INT12 show a severe decrease in MT Polymerase (90%). MT polymerization rates are reported in Table S1 and all values are statistically significantly different from wt-Alp14 to p < 10^−11^. D) Kymographs of individual MT polymerization events with wt or mutants at 200 mM KCl. Top a model for the observed MT polymerization activity and example kymographs shown below. INT1 tracks polymerizing MT-plus ends. INT2 tracks MT plus ends, with short tracking events (white arrows) or rapid unbinding events (red arrows). INT12 shows rapid appearance and disappearance of signal at MT plus ends and slow polymerization rates (red arrows).

Using MT dynamic TIRF assays we studied the MT plus-end tracking and polymerase activities of INT1, INT2, and INT12 mutants in comparison to wt-Alp14 (Table S1; Figure 4C). INT1 shows a 49.1% decrease in maximal MT polymerase (0.96 ± 0.03 μm/min) compared to wt-Alp14. Additionally, INT2 displays a 67.9% loss in maximal MT polymerase activity (0.75 ± 0.02 μm/min). Mutating both interfaces simultaneously, INT12 mutant causes the greatest loss of MT polymerase activity of any Alp14 mutant studied so far *in vitro* (0.53 ± 0.02 μm/min, which is very comparable to the MT polymerization rate without Alp14 (0.38 ± 0.01 μm/min) (Figure 4C; Table S1). These data suggest that despite that the ability of INT1, INT2, and INT12 to bind and recruit αβ-tubulins similarly to wt-Alp14, these mutants lack the square organization, which suggests it serves a critical role in MT polymerase activity.

### TOG square assembly interfaces are essential for processive MT plus-end tracking

The INT1, INT2 and INT12 mutants exhibited abnormal MT plus-end tracking behaviors compared to wt-Alp14 or TOG inactivated mutants. The wt-Alp14 associates with MT plus-ends extensively and resides at polymerizing MT plus-ends for extended periods of time (Figure 1E). In contrast, INT1 tracks the MT-plus ends but displays repeated association and dissociation events, each of which are on 10-40 sec (Figure 4D). INT2 associates with MT-plus ends in short durations, similar to the TOG1M, and also displays similar repeated cycles of association and dissociation during MT polymerization (Figure 4D; red arrows). INT12 displayed a severe defect in MT plus-end tracking, where its signal at MT plus-ends shows a high frequency of association and dissociation events during polymerizing phases, each of which is roughly 10-40 seconds (Figure 4D). We measured the variations in the tracking ratios for INT1, INT2, and INT12, using the approach described in Figure 2. Our analysis shown in Figure 5A-B (additional example kymographs are shown in Figure 5-Supplement 1B-D), reveals that at 200 nM concentrations, INT1 (average: 76.4; range: 84.3%), INT2 (average: 66.4%; range: 97.5%), and INT12 (range: 86.1; average: 78.3%) possess high variation in bulk tracking residence ratios at MT plus-ends compared to wt-Alp14 (range: 36.9%; average: 89.4%) (Figure 5B).

**Figure 5:**
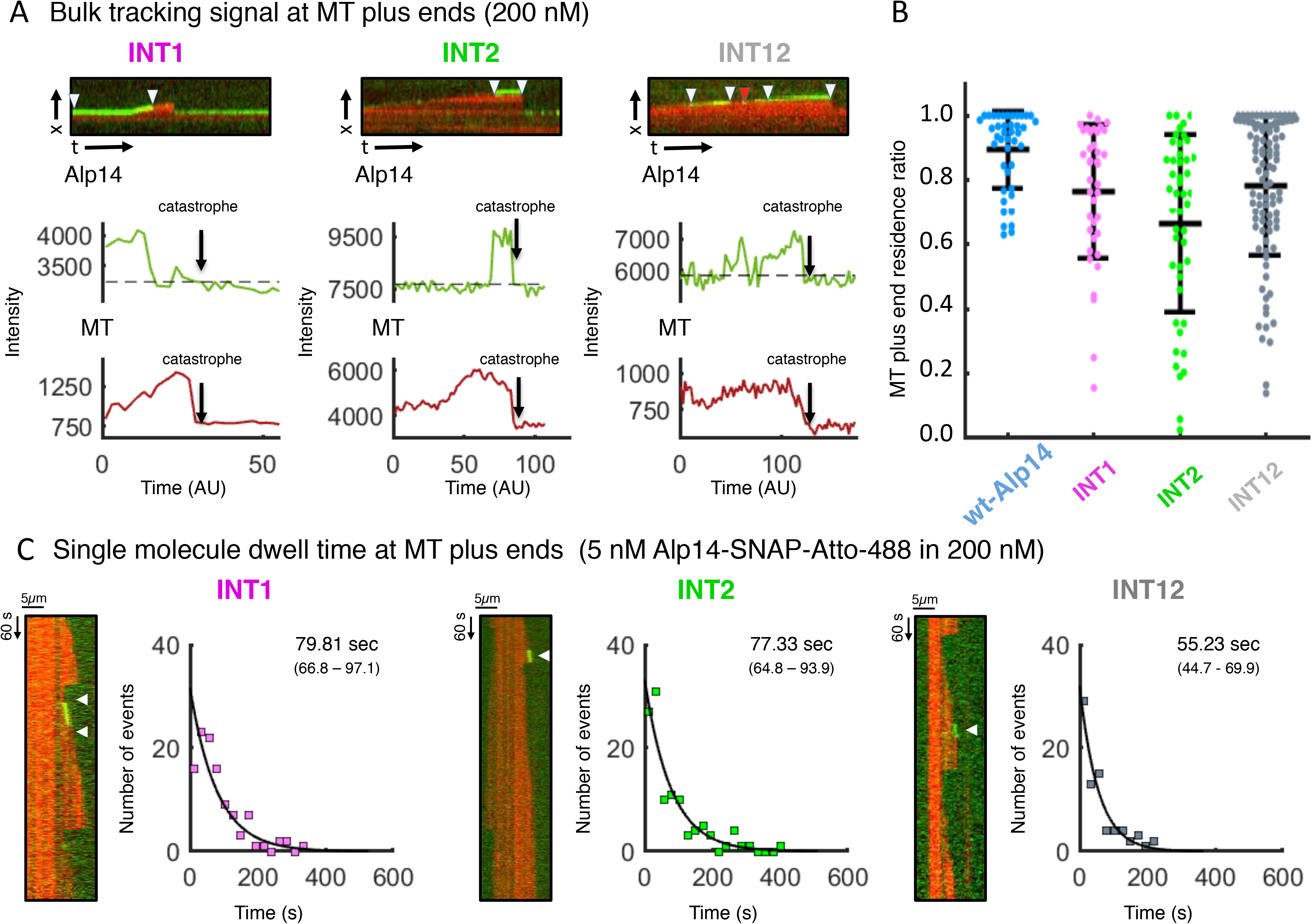
TOG square assembly interfaces are critical for processive MT polymerase activity at MT plus ends. A) Measuring tracking ratio examples for INT1, INT2, and INT12 at MT plus ends. Top panels, horizontal kymograph image, Middle panel, Alp14-bulk intensity that is above background (broken line). Lower panel, MT plus-end intensity throughout their polymerization phase up to catastrophe (arrow) above background (broken line). Note for wt-Alp14 bulk signal fully follows polymerizing MT plus-ends, while INT1, INT2 and INT12 show progressively decreased MT plus end tracking. INT1 shows the mildest zones of no signal denoted by drops below the background in lower panel. Additional examples are shown in Figure 5-Supplement 1B-D. B) MT plus-end residence / persistence Tracking ratios for INT1, INT2, INT12 compared to wt-Alp14. As shown in figure 2, tracking ratios for wt-Alp14 show tight 80-100% distribution. In contrast, tracking ratios show a high variable distribution in INT1, INT2 and INT12 mutants. All values marked with (*) are statistically significant to p < 10^−4^. C) Single molecule dynamic MT polymerization studies reveal the average dwell time for INT1, INT2, and INT12 mutants. Each panel consists of example kymograph (left), dwell time frequencies (right), fit to an exponential decay model. Note that INT1, INT2 and INT12 all show defects in MT plus end tracking. Additional example kymographs are shown in Figure 5-Supplement 2B-E.

To further characterize these MT plus end tracking defects, we measured the MT-plus end dwell times of INT1, INT2, and INT12 at the single molecule level using the approach described in Figure 2C. Individual Atto-488-fluorophore labeled INT1, INT2 and INT12 SNAP-tagged molecule tracking events were identified in kymographs of polymerizing dynamic MTs (Figure 5C; Figure 5-Supplement 2). We measured a large collection of events for each mutant, and used these data to determine the average dwell time at MT plus ends. Interestingly, INT1 and INT2 dwell times along dynamic MT-plus ends are 79.8 sec (n=112) and 77.3 sec (n=112), respectively, which are 45% lower than wt-Alp14 (Figure 5C). The INT12 shows a cumulative defect with a further decrease in dwell time 55.2 sec (n=77), which are roughly 3-fold lower than wt-Alp14. This drop in dwell time in the INT12 mutant suggest that each type of interface is necessary for Alp14 processive tracking at MT plus-ends and the combined defects from the isolated interface lead to a more severe defect when both are inactivated together in the INT12 mutant (Figure 1-Supplement 1B). Overall, the *in vitro* analysis of the interface inactivated mutants suggest that each type of interface plays a different role in organizing arrays of TOG domains which are necessary for proper MT polymerase function of Alp14. Using single molecule dwell times and MT polymerase rates, we calculated the rate of polymerization in αβ-tubulin subunits added per MT-end or protofilament per binding event, as described in Table S2. On average a single wt-Alp14 dimer molecule promotes the polymerization of ∼422 αβ-tubulin subunits per event, INT1 promotes ∼178 αβ-tubulin subunits, INT2 promotes ∼128 αβ-tubulins and INT12 promotes ∼93 αβ-tubulin subunits (Table S2). These data suggest that the square organization is critical for processive Alp14 MT plus-end tracking and thus its inactivation leads to a severe loss of MT polymerase activity in these mutants.

### Inactivating TOG-square interfaces causes defects in tracking of MT plus-ends *in vivo*

We next sought to determine the physiological roles of the TOG square assembly *in vivo*, by introducing these INT1, INT2 and INT12 mutants into *S. pombe* cells. As with the TOG mutants (Figure 3), we replaced the chromosomal Alp14 ORF with INT1, INT2, and INT12 mutants in untagged and GFP-tagged strains. These strains did not display colony growth defects at 20-36 °C temperature range, and in presence of the MT inhibitor MBC (Figure 3-Supplement 1A). These mutants however exhibited significant defects in MT polymerization rates (Figure 6B): INT1 and INT2 showed about a 28% decrease in MT polymerization rate (2.6 ± 0.7 μm/min and 2.4 ± 0.6 μm/min, respectively), while INT12 exhibited a 40% decrease (2.3 ± 0.8 μm/min). This loss of MT polymerase activity resembles the pattern seen in the *in vitro* experiments (Figure 4,5). We also measured similar pattern of decrease in MT depolymerization rates in which INT1, INT2 and INT12 show progressively decreased MT depolymerization rates (Figure 6-Supplement 1B).

**Figure 6:**
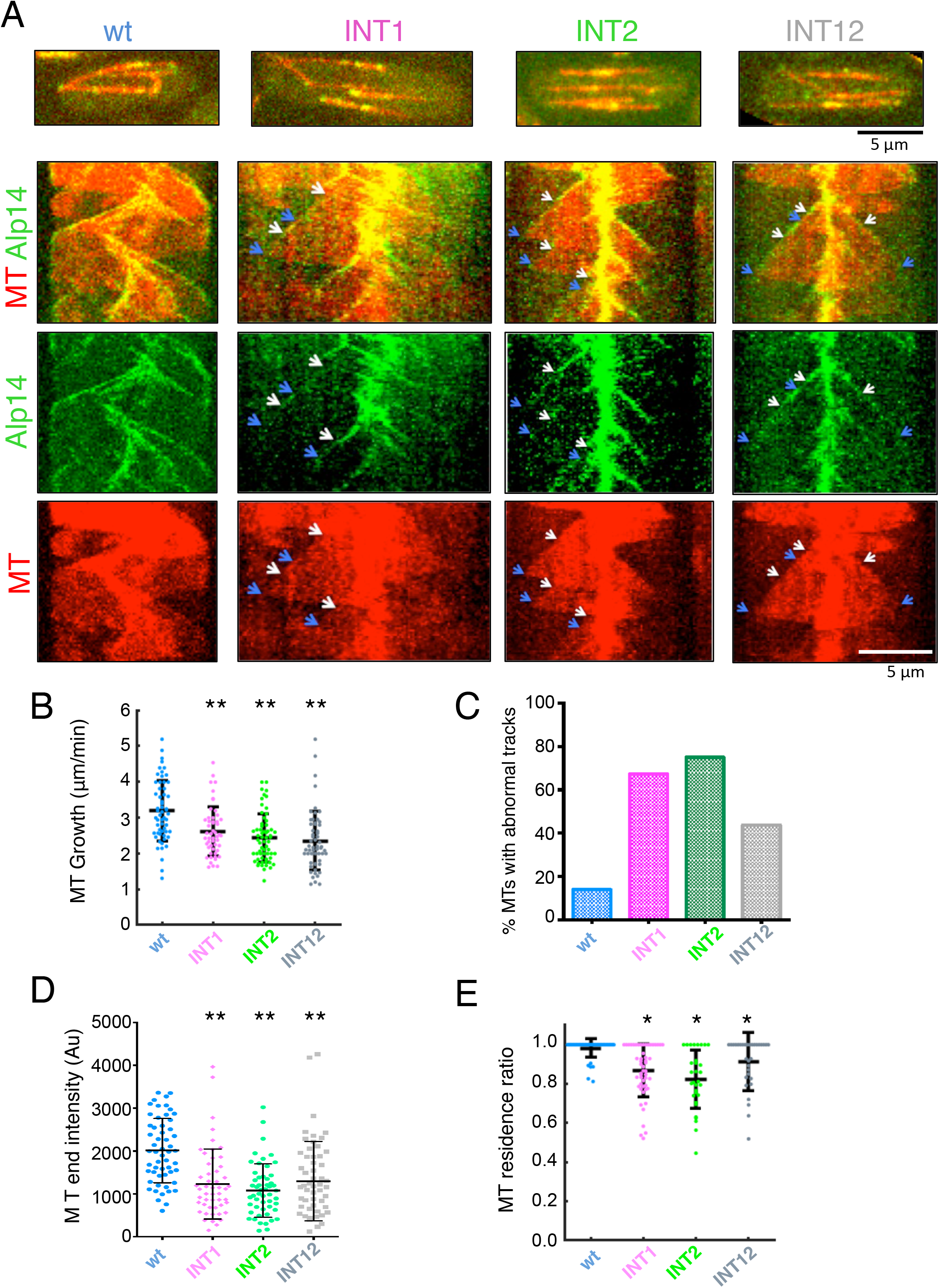
Functional roles for TOG square assembly interfaces in MT polymerase and plus end tracking activities of Alp14 *in vivo*. A) Images of *S. pombe* cells expressing INT1, INT2 and INT12 Alp14 mutants. Top raw cell images with MTs marked with mCherry-Atb2 tubulin (red) and Alp14-GFP (green). Bottom: kymographs of dynamic MTs and Alp14-GFP. Note INT1, INT2, INT12 all track the plus ends of dynamic MTs. White arrows mark the beginning of an abnormal MT tracking event, and blue arrow mark the end of such events. B) MT polymerization rates in wt cells (3.2 um/min), INT1 (2.6 um/min), INT2 (2.4 um/min), INT12 (2.3 um/min) (Table S3). Conditions marked with (**) are statistically significantly different from wt-Alp14 at p < 10^−5^. C) Proportion of dynamic MTs with abnormal tracking behavior in each strain. Note TOG1M, TOG2M and TOG12M have higher ratio of these events. D) Distribution and average of Alp14-GFP intensities at growing MT plus-ends in each of the strains. Note that all the other strains show lower intensity at MT plus-ends E) MT plus-end residence / persistence tracking ratios for Alp14 signal at polymerizing MTs. wt Alp14 shows high tracking ratio (80-100%). INT1 shows a minor decrease in MT plus-end tracking ratio (60-80%), while INT2 and INT12 shows 3-fold decrease in tracking ratio (10-70%). Values are reported in Table S3, and conditions marked with (*) are statistically significantly different from wt-Alp14 at p < 10^−4^.

Like the TOG mutants (Figure 3), the INT mutants showed MT tracking defects (Figure 6A, additional example kymographs are shown in Figure 6-Supplement 1D). Alp14-GFP fluorescence intensities were significantly decreased, with variable distribution of fluorescence intensities at MT plus ends compared to WT cells (Figure 6C). Close visual inspection of time lapse images and kymographs revealed prevalent defects in MT plus end tracking in which Alp14 signals were suddenly decreased for periods on the MT growing end (Figure 6A, additional example kymographs are shown in Figure 6D). Many tracks were dim with uneven intensities (Figure 6A; Figure 6-Supplement 1D). MTs showing some evidence of tracking defects were detected in approximately 40-70% of interphase MTs analyzed (Figure 6D; INT1 67%, INT 75%, and INT12 43% of MTs with abnormal tracking).

Measurement of tracking ratios also revealed MT tracking defects (Figure 6E). INT1 (average: 86.8%; range: 47.9%), INT2 (average: 82.3%; range: 55.4%), and INT12 (average: 91.3%; range: 48.1%) cells show variability in their ability to track the MT plus end compared to WT (average 98.1 %; range 18.7%)(statistically significant differences based on t-test p value of 0.001 for all conditions compared to wt cells). These data support the observations made in the *in vitro* studies (Figure 5) and show that the square assembly of Alp14 TOG arrays are likely to be important *in vivo* for consistent processive tracking at MT plus ends, as well as MT polymerase activity.

## Discussion

### Distinct roles for αβ -tubulin binding and TOG array organization during the MT polymerase cycle

Our *in vitro* and *in vivo* studies suggest that the recruitment of αβ-tubulin by TOG1 and TOG2 domains in arrays is crucial for distinct aspects of MT polymerase and plus-end tracking activities of the XMAP215/Alp14/Stu2 MT polymerases. We also show that non-αβ-tubulin binding interfaces of TOG1 and TOG2 domains, which stabilize the TOG square assembly organization (Nithianantham et al 2018), are also critical for persistent MT plus end tracking leading to defects in MT polymerase activities. Our studies show distinct defects arising from inactivating the αβ-tubulin binding activities for TOG1 and TOG2, compared to inactivating the non-αβ-tubulin binding interfaces 1 and 2 in near native homodimeric Alp14 both *in vitro* and *in vivo*.

The polarized unfurling model suggests that TOG1 and TOG2 domains play distinct roles in MT polymerase and plus-end tracking activities: TOG1 with its high affinity for its αβ-tubulin, anchors the array to the polymerizing MT plus-end, while TOG2 with its rapid exchange for αβ-tubulin, drives αβ-tubulin-αβ-tubulin interaction driving a more rapid polymerization process. However, both TOG1 and TOG2 are critical in the concerted MT polymerase activity. The assembly of a square organization for these arrays is critical in setting the TOG1 and TOG2 positions at polymerizing MT plus ends. The positions of the TOG1 and TOG2 domains are driven by a sequential disassembly of the interfaces stabilizing the square complex upon docking at MT plus end. The specific predictions and defects expected by the polarized unfurling model (Nithianantham et al, 2018) are validated by studies presented here, revealing critical elements in processive MT plus-end tracking, which is a highly conserved feature of XMAP215/Alp14/Stu2 proteins. Our data are consistent with critical roles for multiple TOG domains in Alp14 and XMAP215 MT polymerase activity (Brouhard et al., 2008; Al-Bassam et al, 2012). *In vitro* studies of XMAP215 and Alp14 mutants with inactivated tubulin binding interfaces indicate TOG1 and TOG2 are the most critical for MT polymerase activity, but TOG3 and TOG4 in XMAP215 also play important roles (Al-Bassam et al, 2012; Widlund et al., 2011). Despite their importance for activity, the TOG1 and TOG2 domains are not sufficient as a TOG1-TOG2 construct with charged sequence shows poor activity suggesting full complement of active TOG domains in XMAP215 is essential for MT polymerase activity (Thawani et al., 2018; Widlund et al., 2011).

Our structural studies suggest that two sequential states drive MT polymerase activity: a αβ-tubulin recruitment state and a polymerization-promoting state (Figure 7). The binding of αβ-tubulin by TOG1 and TOG2 domains serve distinct roles in the function of MT polymerases during both recruitment and polymerization states. However TOG square assembly interfaces 1 and 2 only influence formation of the recruitment complex, and their inactivation leads to defects in processive association MT plus ends *in vitro* that may be a cause for the decreased MT polymerase activity. The *in vivo* defects in the INT1, INT2 and INT12 mutants suggest similar loss of persistent MT plus-end tracking. Interface 2 is critical for the transition from the recruitment to polymerization states. Although defects in each aspect of the TOG arrays may decrease in MT polymerase activity, their outcomes lead to distinct functional and localization defects.

**Figure 7:**
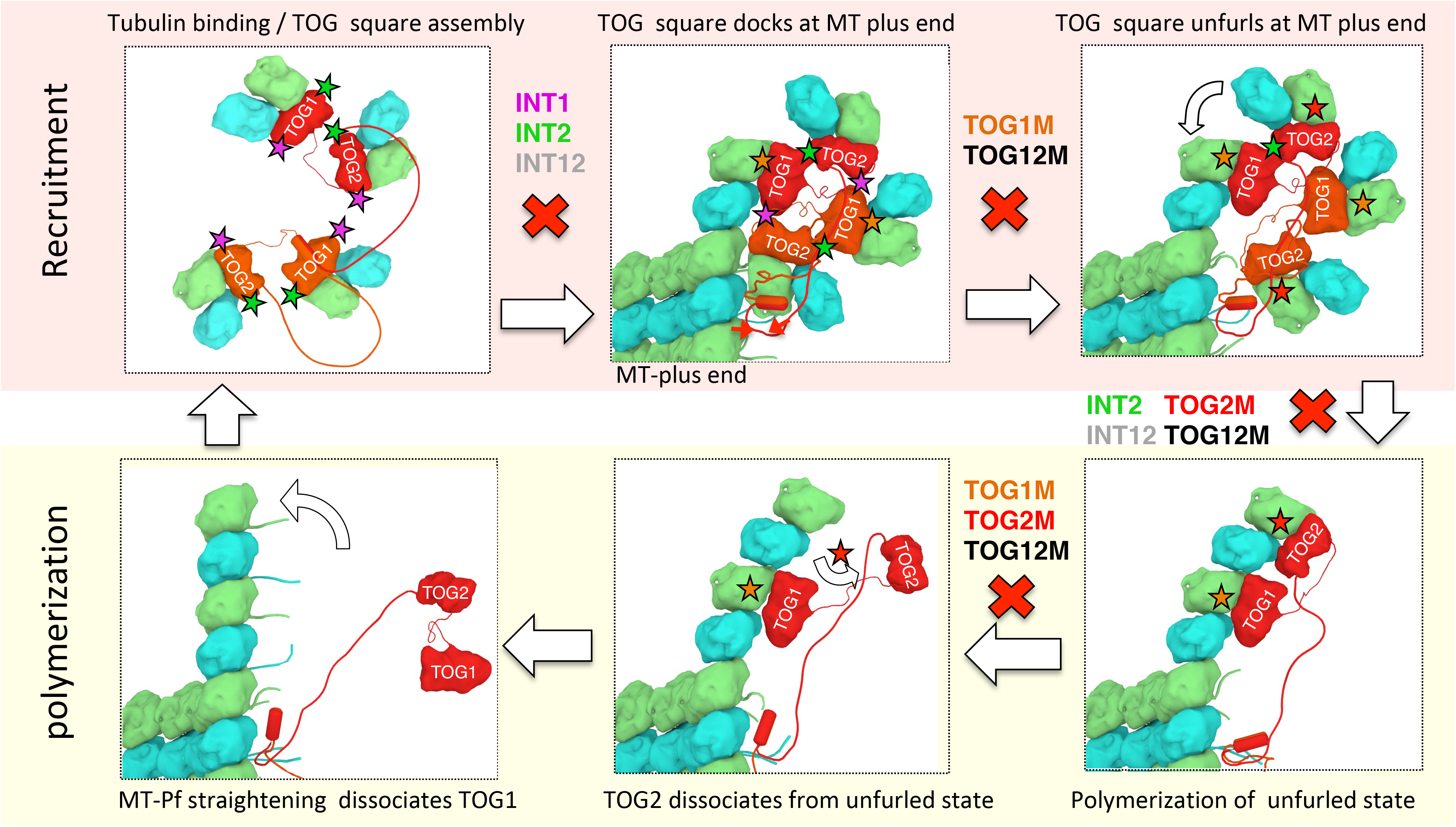
The polarized unfurling model for MT polymerase cycle. Two stages of the MT polymerase polarized unfurling model: Recruitment (red) and Polymerization (yellow). Each includes three stages shown. The transition from each stage to the next may become disrupted (denoted by X) in the mutants are introduced above the transition step. The INT1, INT2, and INT12 mutants disrupt square assembly, and lead to a loss in persistent plus end tracking and decreased polymerization activity. TOG1M, TOG12M mutants, which are deficient in tubulin binding, show defects in persistent plus end tracking, and cannot docking effectively at MT plus ends. and TOG2M and TOG12M are both defective for MT polymerase activity because of a defect in unfurling the TOG square to promote MT polymerization at MT plus ends.

### Molecular basis for processive plus-end tracking by MT polymerases

XMAP215/Stu2/Alp14 family proteins are processive MT plus-end tracking proteins. Alp14 is a typical MT polymerase and includes dimeric arrays of TOG1 and TOG2 domains, followed by a positively charged Ser / Lys rich (Sk-rich) region and dimerization coiled-coil domain. Our studies reveal critical elements for processive MT plus-end tracking by yeast MT polymerases including: TOG1 domain, due to its slow exchange rate of αβ-tubulin, and the square interfaces 1 and 2 that hold together the TOG square organization. Our data indicate that the processive MT plus-end tracking activity is likely due to a cycling from the recruitment and polymerization state. When a complex of Alp14 dimer bound to four αβ-tubulins docks at MT plus ends, TOG1 anchors the assembly and TOG2 is to promote a concerted polymerization of their αβ-tubulins modulated by unfurling the square assembly into a polymerization complex (figure 7). This model fits with observations in the mammalian proteins with their TOG1-TOG5 domains, which likely form a similar compact organization, where TOG1 with its high affinity for αβ-tubulin is critical for MT polymerase activity (Thawani et al., 2018; Widlund et al 2011).

### Comparison to other MT polymerase models

Studies described by other groups suggest alternate models for MT polymerases function. In these models, disorganized arrays of TOG domains simply recruit αβ-tubulins and concentrate them at MT plus-ends and promote lateral αβ-tubulin interactions (Ayaz et al, 2014), or TOG arrays may promote αβ-tubulin assembly into protofilaments in solution prior to docking onto MT plus ends, but with a reverse order in which TOG1 resides at the outer most αβ-tubulin while TOG2 and beyond bind to MTs (Byrnes and Slep, 2017). Both models indicate that recruitment of αβ-tubulins at MT plus-ends is a disorganized process. In contrast to our polarized unfurling model, these models don’t ascribe unique roles to TOG1 and TOG2 domains in the MT plus-end tracking and MT polymerase activities (Rice and Brouhard, 2018). Both of these models suggest only αβ-tubulin recruiting activities, via their αβ-tubulin interfaces, and flexible linkers between TOG domains are critical for MT polymerase activity. These models don’t ascribe any role for square assembly interfaces 1 and 2, which don’t influence αβ-tubulin binding affinity but affect TOG array organization. Our data suggests that the interfaces that hold together the square organization are critical for processive MT plus-end tracking and thus influence MT polymerase activity.

Contrary to previous models, we show the αβ-tubulins recruited by TOG1 and TOG2 do not incrementally add to both the MT polymerase and processive MT plus-end tracking activities. Decreasing αβ-tubulin exchange in TOG2 through decreasing solution ionic strength directly modulates MT polymerase activity. Although both TOG1M and TOG2M show MT polymerase defects, our studies reveal these defects originate from unique MT plus-end tracking activities. TOG1M shows a defect in persistent MT plus-end tracking leading to poor association with MT plus-ends. At 50 mM KCl, TOG1M molecules diffuse away from MT plus ends onto MT lattices (Figure 1). In contrast, TOG2M show no defects in MT-plus end tracking, yet a severe decrease in MT polymerase activity. In both the *in vivo* and *in vitro* settings, the MT polymerase defects observed with TOG2M are consistently more severe than those we observed for TOG1 M (Figure 3). Our studies suggest that MT polymerase and MT-plus end tracking activities are not directly linked to collective activities of TOG domains within the native arrays, but may relate to the different positions of TOG domains during the polymerization process (Figure 7).

The distinct roles of TOG1 and TOG2 domains are consistent with the concerted polymerization state observed in the unfurled polymerized structural model described in the accompanying manuscript (Figure 7; Nithianantham et al 2018). TOG1 bound αβ-tubulin stabilizes the complex at the MT plus-end, while TOG2 drives αβ-tubulin-αβ-tubulin interaction via a rotation and release of its αβ-tubulin on top of TOG1 αβ-tubulin. We tuned the exchange rate by TOG2 in wt-Alp14, either through mutation or manipulating the ionic strength solution conditions, and that modulates the maximal MT polymerase activity without influencing MT-plus end tracking. Our structures suggest that a possible origin of these distinct roles is the asymmetric shape of the TOG square assembly interfaces, which may only become destabilized upon docking of αβ-tubulin bound to TOG1 onto the MT plus end, but not the docking of αβ-tubulin bound to TOG2. As a result, TOG1 becomes the “defacto” site for destabilizing the square complex and the unfurling event (Figure 7).

### TOG square organization is necessary for processive MT plus-end tracking

X-ray structures suggest that Interfaces 1 and 2 stabilize the Alp14 recruitment state (Nithianantham et al 2018). Upon their inactivation, TOG array-αβ-tubulin complexes show either disordered assemblies or spontaneously polymerized assemblies. The *in vitro* and *in vivo* studies here reveal interfaces 1 and 2 are critical for MT plus end tracking and MT polymerase activity despite having no defects in αβ-tubulin association or exchange (Nithianantham et al, 2018). Our *in vitro* studies indicate these mutants display distinct MT polymerase and plus-end tracking defects. INT1 shows an intermediate phenotype with a poor, yet still active, MT polymerase, likely because it still retains the ability form unfurled tubulin intermediates. INT2 shows a severe MT polymerase defect, similar to TOG12M, despite binding αβ-tubulins in a nearly identical manner to wt-Alp14. INT12, which harbors dual inactivation of both interfaces 1 and 2, is fully active in binding αβ-tubulins but shows disordered organization leading to the most severe MT polymerase defect *in vitro*, rapid dissociation defects in tracking MT plus-end and the shortest dwell time at MT plus ends (Figure 4). The *in vivo* the MT polymerase defects of the INT1, INT2 and INT12 mutants (Figure 6) are slight less severe than their MT polymerase defects *in vitro* (Figure 4). The reason for the latter remains unclear, but a cellular factor(s) may stabilize the square assembly conformation leading to a milder defect *in vivo*. The organization of the TOG array thus directly modulates MT plus-end tracking dwell time and controls MT polymerase activity.

### TOG array unfurling via Interface 2 is essential for MT polymerase

Our *in vitro* and *in vivo* studies suggest that interface 2 inactivation leads to a more severe defect than interface 1. This can be explained by the structural transition of TOG square-assembly recruitment state to the unfurled polymerization state (Figure 7). The inter-subunit stabilizing interface 1 may only be critical to stabilize dimeric TOG arrays into TOG squares. A dimerization coil-coil domain maintains the TOG1-TOG2 subunit dimer and thus interface 1 may become dispensable, once Alp14 molecules attach to MT plus ends. Its inactivation in INT1 mutant leads to forming 16-nm polymerized complexes suggesting spontaneous αβ-tubulin polymerization in solution (NIthianantham et al, 2018). In contrast, interface 2 is critical during each polymerization event, which involves an unfurling of TOG2 around TOG1 mediated by Brownian motion (Figure 7). Thus, inactivation interface 2 likely will dramatically increase the diffusion distance of the αβ-tubulins recruited by TOG2 and from αβ-tubulin recruited by TOG1 at the MT plus-end. The similarity of INT2 and TOG1M defects in their poor MT plus-end tracking features indicate the unfurling of TOG2 along a bound TOG1 is likely a critical step for processive tracking and it directly influences MT polymerase activity, if either is inactivated this leads to loss of processive association.

## Materials and Methods

### Cloning, expression and purification of Alp14 and mutants

Coding regions for MT polymerases from *Schizosaccharomyces pombe* Alp14 (accession: BAA84527.1) or its mutants were inserted into a T7 bacterial expression vectors as either a C-terminal Gly-Ser linker (single amino acid code: GSGS) followed by mNeonGreen coding sequence (Allele Biotechnology) and 6Xhis-tag, or a C-terminal Gly-Ser linker followed by SNAPf coding sequence (NEB #N9183S) and his-tag (Figure 1 Supplement 1A). These constructs included residues 1-690 of Alp14 including TOG1, TOG2 and the Ser-Lys rich (Sk-rich) region and coiled-coil dimerization domains (Figure 1 Supplement 1A). To generate TOG1M and TOG2M mutants, point mutagenesis was used to introduce Trp23Ala, Arg109Ala, and Arg200Ala to inactivate TOG1 (TOG1M) and Trp300Ala, Lys381Ala, and Arg463Ala to inactivate TOG2 (TOG2M) domains and those were combined to generate TOG12M as described in the accompanying manuscript. Gene synthesis was used to generate the INT12 mutant (Epoch life sciences). INT1 was generated by placing a segment of the INT12 ORF containing INT1 mutations (junction TOG1-TOG2 linker) into wt-Alp14 ORF. INT2 was generated by the INT12 ORF segment containing INT2 (outer N and C-terminal edges of TOG1 and TOG2 domains, respectively) were placed in wt-Alp14. All constructs were fully sequenced to confirm the detailed mutations. All constructs were transformed and expressed in SolubBL21 bacterial strains using the T7 expression system, and were grown at 37°C and induced with 0.5 mM isopropyl thio-β-glycoside (RPI) at 18 C overnight. For purification, cells were centrifuged, resuspended in lysis Buffer (50mM HEPES, 400mM KCl, 5mM 2-MercaptoEthano (BME), pH6.8), and then lysed using a microfluidizer (Avastin). Bacterial extracts were clarified via centrifugation at 18,000 x g. Supernatant was passed through 10ml Ni-IDA resin (Macherey-Nagel) and eluted using Elution Buffer (50mM HEPES, 400mM KCl, 5mM BME, 250mM Imidazole). Elution fractions was then diluted in Salt Cut buffer (50mM HEPES, 50mM KCl, 5mM BME, pH6.8) to approximately 200mM KCl buffer concentration and loaded onto 10ml ion exchange Hitrap-SP (GE) chromatography. The protein was then eluted using a 20 column volumes (CVs) in linear gradient between a Low salt buffer (50mM HEPES, 200-500 mM KCl, 5mM BME, pH6.8) and a High salt Buffer (50mM HEPES, 500mM KCl, 5mM BME, pH6.8). Elution fractions were pooled and concentrated using a 50,000 MWCO Amicon Ultra Centrifugal Filter (Millipore Sigma) and loaded onto Size exclusion gel-filtration chromatography using a Superdex 200 (30/1000) column (GE Healthcare). Purified proteins were used immediately for a period of 3-4 days for dynamic TIRF assays.

### In vitro dynamic MT polymerization assays

MT polymerase activity with dynamic MTs was reconstituted using Total internal microscopy was carried out as described previously (Al-Bassam, 2014) with the following important modifications: Flow chambers were assembled from N 1.5 glass coverslips (0.16 mm thick; Ted Pella) which were cleaned with the Piranha protocol and then functionalized with 2 mg/ml PEG-2000-Silane containing 2 ug/ml biotin-PEG-3400-Silane (Laysan Bio) suspended in 80% at pH 1 (Henty-Ridilla et al., 2016). After flow chamber assembly, 0.1mg/ml NeutrAvidin (Thermofisher) was used to functionalize surfaces. Biotin and Alexa-Fluor-633 labeled porcine tubulin were polymerized using the non-hydrolyzable GTP analog GMPCPP (Jena biosciences). 100 ug/ml MT seeds were flowed into the chambers. Dynamic MT polymerization was reconstituted at the 37 °C by injecting into flow chambers a mixture of 8 μM soluble αβ-tubulin containing 5-10% Alexa-Fluor-546-labeled or Alexa-Fluor-633-labeled αβ-tubulin, in the presence of Alp14-NG or Alp14-SNAP proteins or their mutants in BRB-80 (80 mM PIPES, 1 mM MgCl2, 1 mM EGTA, pH 6.8) in 50-200 mM KCl, combined with a photobleach correction mixture (Telley et al., 2011). Movies were captured in TIRF mode using a Nikon Eclipse Ti microscope using 1.5 Na objective using an Andor IXon3 EM-CCD operating with emission filters using alternating filter wheel in 2 sec increments.

### Analysis of MT dynamic polymerization parameters

The movies were preprocessed by using photobleach correction and image stabilization plugins using the program FIJI (Schindelin et al., 2012). Dynamic MTs were identified and kymographs were generated for multiple channels. Rates of MT polymerization, depolymerization and their times were determined from individual Kymograph measurements as well as residence of Alp14 molecules at MT plus ends (Al-Bassam et al, 2010; Al-Bassam, 2014). Large collections of dynamic events for Alp14 or each of its mutants were collected for a variety of solution conditions (Table S1). Average MT parameters were determined from Gaussian distributions using MATLAB (Table S1). In general all parameters had a single Gaussian distribution. Comparison of parameters shown in Figure 1 and 3 were determined by fits using MATLAB using the following formula.

### Analysis of MT plus end Alp14 residence / persistence ratio

Kymographs of dynamic MT plus ends with 200 nM Alp14-NG or its mutants were used to perform line scans of the MT plus ends in the 633 nm and 488 emission channels. Figure 2 Supplement 1 shows how such line scans were performed. Generally Alp14-NG bulk signal was analyzed during MT polymerization event until catastrophe initiation and included a background measurement. As shown in Figure 2 Supplement 1, the 633 nm channel was used to determine the boundary for MT catastrophe and then the boundary was used in the 488 nm channel to identify the Alp14-NG signal through comparison to background. The level of background signal was then used to create a cutoff in the 488 nm channels for Alp14-NG signal, and any signal above this background cutoff was then counted to determine the duration of Alp14-NG tracking. A variety of events for Alp14-NG and each of its mutants are shown in Figure 2 Supplement 1. To determine the residence ratio, the length of Alp14-NG intensity events above background was then compared to the total length of MT growth events, which represents MT growth time. For each condition, 30-50 events were then analyzed and then compared as shown in figure 2 and 4. Similar analyses were used for *in vivo* determination of residence ratios.

### Single molecule TIRF assays to determine Alp14 dwell time at MT plus ends

SNAPf-tagged wt-Alp14 (termed Alp14-SNAP) or its mutants were purified, as described for Alp14-NG and as shown in Figure 1 Supplement 1. 10 uM SNAP-Surface Atto 488 (also named Benzyl-guanine –Atto-488; NEB) was incubated with 5 uM Alp14-SNAP at 4 °C in 50 mM HEPES, 400 mM KCl, 1 mM DTT. Excess SNAP-Atto-488 was removed using Zebra spin desalting columns (Thermofisher). Flow chambers functionalized with MT seeds as described above were injected with mixture containing 5 nM Atto-488 labeled Alp14-SNAP, unlabeled 195 nM Alp14-SNAP and 8 uM tubulin containing 5-10% Alexa-Fluor-633 and then imaged with the system described above. The laser power conditions and fluorphore concentrations studied to determine photo-bleaching profiles and confirm that no photobleaching occurred during the time of the experiment.

Movies containing single Alp14-SNAP MT plus-end tracking events, as seen in Figure 1 Supplement 2, were processed as described above. Individual Alp14-SNAPf events were distinguished by an immediate rise in the signal and the similar levels of signal observed above background. The low ratio (2.5%) of labeled Alp14-SNAPf molecules used in our experiments, ensured that single binding/dissociation events did not overlap and we observed single events in our assays. The frequency of durations for 100-150 events for wt-Alp14-SNAPf were counted and fit to an exponential decay model to determine the t (1/2) for Alp14 dwell time on the MT end.

### Generation of *S. pombe* strains with *alp14* mutants for functional studies

*S. pombe* strains are described in Table S4 and were generated by replacing mutant *alp14* alleles into the endogenous Alp14 locus. Mutants were made as follows: N-terminal Alp14 sequence was amplified from the T7 bacterial expression vectors (described above and in the accompanying manuscript) using the primers ONF89 and ONF91. C-terminal Alp14-GFP-KanMX sequence was amplified from strain FC1907 (Al-Bassam et al, 2012) with primers ONF90 and ONF92 generating an overlapping sequence of 25 nucleotides. Both fragments were ligated using the NEBuilder HiFi Assembly and following manufacture instructions. The resulting amplicon was transformed in *alp14*Δ *S. pombe* cells, confirmed by PCR and sequencing, and outcrossed with wt strain three times. Crosses and tetrad analyses were used to generate double mutants. For sensitivity drop assays, strains were grown in YES cultures to exponential midlog phase at 25°C. Serial 5-fold dilutions were made, starting from a O.D. of 0.2 and spotted onto YES agar plates. For MBC sensitivity assays, cells were spotted onto YES containing DMSO or YES plates containing the indicated concentrations of MBC (Sigma-Aldrich) and grown at 20°C, 25°C, 30°C and 36°C. Pictures were taken after incubation for 3-6 days.

### Imaging and Analysis of Alp14 and MTs in *S. pombe* Cells

For live cell imaging, cells were typically grown in YES at 25 °C for 36 hours keeping them at a density below 0.5 OD_600_. Cells were placed on microslide wells (Ibidi #80821) coated with soybean lectin (Sigma #1395). Images were acquired using a spinning-disk confocal microscope (IX-81; Olympus; CoolSnap HQ2 camera, Plan Apochromat 100×, 1.4 NA objective [Roper Scientific]). Temperature was stably controlled in the room during imaging at 25 °C unless otherwise indicated.

Time-lapse sequences were taken every 4 seconds with 13 Z-sections of 0.5 μm for a total time of 3-5 min. Image stacks were analyzed using ImageJ (Schneider et al., 2012). Maximum intensity Z-projections were used for generating kymographs and intensity analysis. Kymographs and movies of Alp14-GFP and mCherry-Atb2 were used to determine MT growth and shrinkage rates and tracking behavior. The analyses were restricted to growing plus ends of “lead” MTs that grow out towards the cell tips, as these are likely to be single MTs (approach is described in Figure 3-Supplement 2A-C) (Hoogs et al., 2007). MT growth and shrinkage measurements were derived from analyses of kymographs. Alp14-GFP intensities at growing plus ends of lead MTs were measured from single images by mean intensity within a 12-pixel region of interest (ROI) and subtracting the cytoplasmic background intensity from a nearby ROI. In addition to the background subtraction, the “Find Edges” plugin (Process/Find Edges) using FIJI imageJ provided sharper image regarding GFP intensity in the kymograph (Figure 3-Supplement 2C); these GFP dots were confirmed to be growing MT plus ends by examination of time lapse sequences of the same cells (Figure 3-Supplement 2D). Alp14-GFP tracking of MTs was evaluated visually on each individual MT by comparing the GFP signal with the behavior of the MT on kymographs and corresponding movie frames. MTs ends that showed sudden loss of GFP signal (very dim or absent signal on the plus end) were scored as having abnormal tracking (Figure 3D,6D). The measurements of residence ratios were determined in a similar manner as described for *in vitro* analyses, but include periods of dim but detectable signal as well as undetectable signal (Figure 3E,6E).

## Statistical Analysis

Graphs and statistical analyses were performed using GraphPad Prism version 5.0 (GraphPad Software).

---

**Table S1:** MT dynamic polymerization parameters reveal activities of Alp14 and mutants as measured by TIRF assays

| TIRF Conditions | Growth rate<br>(um/min) | Growth time (sec) | Shrinkage rate<br>(um/min) | Shrinkage time<br>(min) | catastrophe<br>frequency | MT length<br>(um) | MT Residence /<br>Persistence Ratio |
| --- | --- | --- | --- | --- | --- | --- | --- |
| 8uM Tubulin, 200mM KCl | 0.38±.009 n=132 | 383.±21.5 n=132 | 40.8±1.6 n=95 | 0.077±.003 n=95 | 0.28±.02 n=96 | 2.533±.164 n=132 | N/D |
| 50nM wt-Alp14, 200mM KCl | 1.042±.031 n=121 | 132.1±6.8 n=121 | 54.6±2.8 n=82 | 0.069±.003 n=82 | 0.831±.065 n=83 | 2.366±.15 n=121 | N/D |
| 100nM wt-Alp14, 200 mM KCl | 1.363±.031 n=110 | 141.9±7.3 n=110 | 50.7±2.8 n=72 | 0.081±.003 n=72 | 0.699±.043 n=72 | 3.192±.172 n=110 | N/D |
| 150nM wt-Alp14, 200 mM KCl | 1.455±.027 n=99 | 147.5±7.8 n=99 | 56.6±4.4 n=71 | 0.091±.004 n=71 | 0.569±.036 n=72 | 3.486±.18 n=99 | N/D |
| 200nM wt-Alp14, 200 mM KCl | 1.524±.033 n=146 | 123.5±5.6 n=146 | 63.7±2.6 n=95 | 0.074±.003 n=95 | 0.77±.049 n=101 | 3.191±.16 n=146 | 89.42 ± 12.0 n= 43 |
| 50nM TOG1M, 200 mM KCl | 0.758±.027 n=103 | 156.3±8.9 n=103 | 53.±2.9 n=80 | 0.06±.003 n=80 | 0.559±.038 n=84 | 1.919±.116 n=103 | N/D |
| 100nM TOG1M, 200 mM KCl | 0.889±.025 n=96 | 114.4±7.1 n=96 | 49.1±2.7 n=70 | 0.061±.003 n=70 | 0.796±.058 n=76 | 1.777±.128 n=96 | N/D |
| 150nM TOG1M, 200 mM KCl | 0.878±.02 n=121 | 100.1±5.7 n=121 | 54.9±2.7 n=93 | 0.054±.002 n=93 | 0.973±.068 n=97 | 1.514±.097 n=121 | N/D |
| 200nM TOG1M, 200 mM KCl | 0.834±.025 n=115 | 92.1±4.8 n=115 | 44.9±2.3 n=87 | 0.056±.003 n=87 | 0.937±.053 n=87 | 1.33±.088 n=115 | 72.38 ± 23.3 n= 32 |
| 50nM TOG2M, 200mM KCl | 0.744±.026 n=91 | 117.7±7.3 n=91 | 43.5±2.5 n=68 | 0.053±.002 n=68 | 0.724±.053 n=70 | 1.458±.099 n=91 | N/D |
| 100nM TOG2M, 200 mM KCl | 0.83±.02 n=138 | 123.±6.4 n=138 | 45.9±2.1 n=111 | 0.05±.002 n=111 | 0.761±.039 n=115 | 1.741±.104 n=138 | N/D |
| 150nM TOG2M, 200 mM KCl | 0.652±.018 n=121 | 95.6±4.6 n=121 | 35.9±1.9 n=98 | 0.051±.002 n=98 | 0.901±.051 n=105 | 1.047±.061 n=121 | N/D |
| 200nM TOG2M, 200 mM KCl | 0.701±.02 n=121 | 91.5±5.4 n=121 | 34.5±1.6 n=89 | 0.049±.002 n=89 | 1.052±.069 n=107 | 1.054±.07 n=121 | 91.09 ± 11.1 n= 34 |
| 50nM TOG12M, 200 mM KCl | 0.664±.023 n=82 | 192.9±11.7 n=82 | 64.5±3.4 n=50 | 0.054±.003 n=50 | 0.525±.044 n=58 | 2.309±.188 n=82 | N/D |
| 100nM TOG12M, 200 mM KCl | 0.869±.035 n=58 | 201.4±15.2 n=59 | 72.6±5.5 n=35 | 0.06±.003 n=35 | 0.499±.053 n=37 | 3.013±.31 n=59 | N/D |
| 150nM TOG12M, 200 mM KCl | 0.753±.019 n=90 | 236.2±12.4 n=90 | 70.9±4.3 n=54 | 0.066±.003 n=54 | 0.431±.041 n=57 | 3.045±.183 n=90 | N/D |
| 200nM TOG12M, 200 mM KCl | 0.648±.016 n=101 | 183.3±8.8 n=101 | 56.3±2.7 n=79 | 0.068±.003 n=79 | 0.496±.037 n=80 | 2.011±.112 n=101 | N/D |
| 50nM INT1, 200 mM KCl | 0.879±.021 n=146 | 112.5±5.1 n=146 | 29.8±1. n=110 | 0.08±.004 n=110 | 0.784±.05 n=114 | 1.618±.079 n=146 | N/D |
| 100nM INT1, 200 mM KCl | 0.999±.022 n=144 | 112.3±5.6 n=144 | 47.3±1.9 n=113 | 0.05±.002 n=113 | 0.788±.049 n=123 | 1.809±.088 n=144 | N/D |
| 150nM INT1, 200 mM KCl | 0.953±.028 n=124 | 103.3±5.8 n=124 | 52.5±2.5 n=92 | 0.043±.001 n=92 | 0.859±.045 n=103 | 1.584±.092 n=75 | N/D |
| 200nM INT1, 200 mM KCl | 0.957±.03 n=102 | 104.4±5.6 n=101 | 52.±2.9 n=83 | 0.052±.002 n=83 | 0.85±.066 n=88 | 1.672±.111 n=101 | 76.41 ± 20.9 n= 38 |
| 50nM INT2, 200 mM KCl | 0.672±.014 n=168 | 204.8±9.6 n=167 | 65.3±2.2 n=104 | 0.057±.01 n=104 | 0.475±.025 n=119 | 2.476±.134 n=154 | N/D |
| 100nM INT2, 200 mM KCl | 0.69±.016 n=127 | 225.1±12.2 n=127 | 45.9±1.8 n=82 | 0.08±.004 n=82 | 0.428±.027 n=84 | 2.597±.149 n=127 | N/D |
| 150nM INT2, 200 mM KCl | 0.73±.021 n=136 | 227.±10.8 n=136 | 60.3±2.3 n=82 | 0.075±.004 n=82 | 0.465±.044 n=86 | 2.76±.151 n=136 | N/D |
| 200nM INT2, 200 mM KCl | 0.747±.018 n=173 | 167.7±6.7 n=172 | 57.±1.8 n=122 | 0.063±.01 n=122 | 0.507±.029 n=134 | 2.116±.102 n=156 | 66.41 ± 27.5 n= 49 |
| 50nM INT12, 200 mM KCl | 0.488±.015 n=129 | 72.±5.7 n=129 | 37.7±2. n=96 | 0.023±.001 n=96 | 1.935±.16 n=104 | .454±.039 n=100 | N/D |
| 100nM INT12, 200 mM KCl | 0.473±.015 n=108 | 134.5±6. n=108 | 33.±1.6 n=65 | 0.05±.002 n=65 | 0.631±.033 n=77 | 1.065±.06 n=100 | N/D |
| 150nM INT12, 200 mM KCl | 0.579±.018 n=92 | 135.1±6.9 n=92 | 51.4±2.6 n=61 | 0.045±.002 n=61 | 0.604±.042 n=69 | 1.366±.091 n=92 | N/D |
| 200nM INT12, 200 mM KCl | 0.526±.017 n=128 | 83.3±5.5 n=128 | 44.8±2.1 n=93 | 0.029±.001 n=93 | 1.436±.122 n=115 | .719±.055 n=128 | 78.35 ± 21.7 n= 97 |
| 8uM Tubulin, 50mM KCl | 0.499±.012 n=108 | 293.5±17.7 n=108 | 30.3±1.3 n=69 | 0.128±.008 n=69 | 0.423±.039 n=70 | 2.528±.179 n=108 | N/D |
| 50nM wt-Alp14, 50mM KCl | 1.042±.023 n=181 | 121.4±5.8 n=181 | 32.9±1.6 n=126 | 0.099±.01 n=127 | 0.902±.056 n=132 | 2.133±.112 n=181 | N/D |
| 100nM wt-Alp14, 50mM KCl | 1.016±.148 n=150 | 114.±5.8 n=150 | 41.1±1.9 n=107 | 0.122±.02 n=109 | 0.845±.056 n=109 | 2.266±.143 n=150 | N/D |
| 150nM wt-Alp14, 50mM KCl | 1.15±.026 n=127 | 114.2±5.4 n=127 | 37.3±1.7 n=92 | 0.08±.004 n=92 | 0.783±.043 n=93 | 2.24±.126 n=127 | N/D |
| 200nM wt-Alp14, 50mM KCl | 1.073±.037 n=128 | 128.8±6. n=128 | 39.4±2. n=87 | 0.097±.005 n=87 | 0.666±.04 n=87 | 2.302±.132 n=120 | N/D |
| 50nM TOG1M, 50mM KCl | .741±.021 n=105 | 234.3±12.6 n=105 | 17.5±.8 n=74 | 0.34±.027 n=74 | 0.409±.037 n=74 | 2.869±.213 n=52 | N/D |
| 50nM TOG2M, 50mM KCl | .328±.011 n=102 | 225.4±12.6 n=107 | 13.5±.7 n=73 | 0.176±.013 n=73 | 0.452±.032 n=78 | 1.887±.257 n=72 | N/D |

**Table S2:** Dwell times and subunits polymerized for Alp14 mutants at MT plus ends

| Protein | Dwell time (sec), 95% confidence interval | MT Growth Rate (subunit / sec )<sup>a,b</sup> | $\alpha\beta$-tubulins added to 13-pf MT per event: | $\alpha\beta$-tubulins added to 1 pf per event<sup>a,c</sup> |
| --- | --- | --- | --- | --- |
| **wt-Alp14** | 129.29 n=164 (111.43 - 151.84) | 42.52 | 5497.52 | 422.89 |
| **TOG1M** | 57.45 n=130 (48.73 - 68.77) | 22.56 | 1296.10 | 99.7 |
| **TOG2M** | 132.37 n=152 (113.61 - 156.22) | 18.99 | 2511.58 | 193.2 |
| **INT1** | 79.82 n=112 (66.77 - 97.12) | 29.11 | 2323.66 | 178.74 |
| **INT2** | 77.34 n=112 (64.8 - 93.93) | 20.2 | 1562.39 | 120.18 |
| **INT12** | 55.24 n=77 (44.71 - 69.99) | 21.99 | 1214.60 | 93.43 |

a) Calculating dimer addition to protofilament end was calculated as following based on (Brouhard et al, 2008). pf = protofilaments
b) $MT\ GrowthRate\left(\frac{\mu m}{min}\right) * \frac{1000nm}{60sec} * \frac{1\ Dimer}{8nm} = MT\ GrowthRate\left(\frac{dimers}{sec}\right) for\ 13\ pf$
c) $MT\ GrowthRate\left(\frac{dimers}{sec}\right) for\ 13\ pf * Residence\ Time(sec) * \frac{1}{13\ pf} = \# of\ Dimers\ added\ to\ 1\ pf$

**Table S3:** In vivo MT dynamic polymerization / depolymerization parameters and Alp14 intensity and tracking parameters

| ***S. pombe***<br>**strains**<br>**studied** | **MT Growth rate**<br>**($\mu\text{m}/\text{min}$)** | **MT shrinkage rate**<br>**($\mu\text{m}/\text{min}$)** | **MT length**<br>**($\mu\text{m}$)** | **Intensity**<br>**(AU)** | **MT residence/**<br>**persistence ratio** |
| --- | --- | --- | --- | --- | --- |
| WT | 3.188 ± .845 n=58 | 12.088 ± 3.615 n=58 | 6.025 ± 1.764 n=112 | 2012.3 ± 749.9 n=55 | 0.982 ± 0.05 n= 43 |
| TOG1M | 1.557 ± .487 n=42 ** | 10.574 ± 2.636 n=42 ** | 4.822 ± 1.768 n=118 | 1094 ± 6815 n=56 | 0.842 ± 0.17 n= 54 * |
| TOG2M | 1.329 ± .308 n=43 ** | 7.088 ± 1.997 n=43 ** | 4.371 ± 1.49 n=117 | 1419.2 ± 672.2 n=52 | 0.961 ± 0.06 n= 28 <sup>ns</sup> |
| TOG12M | 1.155 ± .25 n=32 ** | 6.881 ± 1.408 n=32 ** | 3.454 ± 1.587 n=117 | 1157.0 ± 885.6 n=57 | 0.564 ± 0.26 n= 15 * |
| INT1 | 2.615 ± .684 n=51 ** | 12.305 ± 4.115 n=51 ** | 5.833 ± 2.232 n=127 | 1230.0 ± 816.1 n=55 | 0.868 ± 0.13 n= 52 * |
| INT2 | 2.427 ± .665 n=63 ** | 10.822 ± 2.522 n=63 ** | 6.042 ± 1.969 n=127 | 1076 ± 622.8 n=53 | 0.823 ± 0.15 n= 31 * |
| INT12 | 2.345 ± .821 n=57 ** | 9.105 ± 2.625 n=57 ** | 5.747 ± 1.879 n=119 | 1298 ± 928.2 N=56 | 0.913 ± 0.13 n= 32 * |

\* marks differences with wt that are significant using student's t-test to the level of $p < 0.001$
\*\* marks differences with wt that are significant using the student t-test to the level of $< 0.0001$
<sup>ns</sup> marks differences that are not significantly different from wt based on the students' t-test.

**Table S4:** genotypes of fission yeast strains used in this study

| Strain # | Genotype | Refs |
| --- | --- | --- |
| NF1241 | *Alp14-GFP:KanMX ura4D18 leu2-11 ade6-M210* | This study |
| NF1247 | *Alp14TOG1-GFP:KanMX ura4D18 leu2-11 ade6-M210* | This study |
| NF1253 | *Alp14TOG2-GFP:KanMX ura4D18 leu2-11 ade6-M210* | This study |
| NF1259 | *Alp14INT1-GFP:KanMX ura4D18 leu2-11 ade6-M210* | This study |
| NF1265 | *Alp14INT2-GFP:KanMX ura4D18 leu2-11 ade6-M210* | This study |
| NF1269 | *Alp14INT12-GFP:KanMX ura4D18 leu2-11 ade6-M210* | This study |
| NF1270 | *Alp14TOG12-GFP:KanMX ura4D18 leu2-11 ade6-M210* | This study |
| NF1416 | *Alp14-GFP:KanMX mCherry-atb2:HphMX ura4D18 leu2-11 ade6-M210* | This study |
| NF1418 | *Alp14TOG1-GFP:KanMX mCherry-atb2:HphMX ura4D18 leu2-11 ade6-M210* | This study |
| NF1420 | *Alp14TOG2-GFP:KanMX mCherry-atb2:HphMX ura4D18 leu2-11 ade6-M210* | This study |
| NF1422 | *Alp14INT1-GFP:KanMX mCherry-atb2:HphMX ura4D18 leu2-11 ade6-M210* | This study |
| NF1424 | *Alp14INT2-GFP:KanMX mCherry-atb2:HphMX ura4D18 leu2-11 ade6-M210* | This study |
| NF1426 | *Alp14INT12-GFP:KanMX mCherry-atb2:HphMX ura4D18 leu2-11 ade6-M210* | This study |
| NF1428 | *Alp14TOG12-GFP:KanMX mCherry-atb2:HphMX ura4D18 leu2-11 ade6-M210* | This study |

## References

Akhmanova, A., and Steinmetz, M.O. (2011). Microtubule End Binding: EBs Sense the Guanine Nucleotide State. Curr Biol 21, R283–285.

Akhmanova, A., and Steinmetz, M.O. (2015). Control of microtubule organization and dynamics: two ends in the limelight. Nat Rev Mol Cell Biol.

Al-Bassam, J. (2014). Reconstituting dynamic microtubule polymerization regulation by TOG domain proteins. Methods Enzymol 540, 131–148.

Al-Bassam, J., and Chang, F. (2011). Regulation of microtubule dynamics by TOG-domain proteins XmAp215/Dis1 and CLASP. Trends Cell Biol 21, 604–614.

Al-Bassam, J., Kim, H., Flor-Parra, I., Lal, N., Velji, H., and Chang, F. (2012). Fission yeast Alp14 is a dose-dependent plus end-tracking microtubule polymerase. Mol Biol Cell 23, 2878–2890.

Al-Bassam, J., van Breugel, M., Harrison, S.C., and Hyman, A. (2006). Stu2p binds tubulin and undergoes an open-to-closed conformational change. J Cell Biol 172, 1009–1022.

Ayaz, P., Munyoki, S., Geyer, E.A., Piedra, F.A., Vu, E.S., Bromberg, R., Otwinowski, Z., Grishin, N.V., Brautigam, C.A., and Rice, L.M. (2014). A tethered delivery mechanism explains the catalytic action of a microtubule polymerase. Elife 3, e03069.

Ayaz, P., Ye, X., Huddleston, P., Brautigam, C.A., and Rice, L.M. (2012). A TOG:alphabeta-tubulin complex structure reveals conformation-based mechanisms for a microtubule polymerase. Science 337, 857–860.

Brouhard, G.J., and Rice, L.M. (2014). The contribution of alphabeta-tubulin curvature to microtubule dynamics. J Cell Biol 207, 323–334.

Brouhard, G.J., Stear, J.H., Noetzel, T.L., Al-Bassam, J., Kinoshita, K., Harrison, S.C., Howard, J., and Hyman, A.A. (2008). XMAP215 is a processive microtubule polymerase. Cell 132, 79–88.

Byrnes, A.E., and Slep, K.C. (2017). TOG-tubulin binding specificity promotes microtubule dynamics and mitotic spindle formation. J Cell Biol 216, 1641–1657.

Flor-Parra, I., Iglesias-Romero, A.B., and Chang, F. (2018). The XMAP215 Ortholog Alp14 Promotes Microtubule Nucleation in Fission Yeast. Curr Biol.

Garcia, M.A., Vardy, L., Koonrugsa, N., and Toda, T. (2001). Fission yeast ch-TOG/XMAP215 homologue Alp14 connects mitotic spindles with the kinetochore and is a component of the Mad2-dependent spindle checkpoint. EMBO J 20, 3389–3401.

Haase, K.P., Fox, J.C., Byrnes, A.E., Adikes, R.C., Speed, S.K., Haase, J., Friedman, B., Cook, D.M., Bloom, K., Rusan, N.M., et al. (2018). Stu2 uses a 15-nm parallel coiled coil for kinetochore localization and concomitant regulation of the mitotic spindle. Mol Biol Cell 29, 285–294.

Henty-Ridilla, J.L., Rankova, A., Eskin, J.A., Kenny, K., and Goode, B.L. (2016). Accelerated actin filament polymerization from microtubule plus ends. Science 352, 1004–1009.

Kakui, Y., Sato, M., Okada, N., Toda, T., and Yamamoto, M. (2013). Microtubules and Alp7-Alp14 (TACC-TOG) reposition chromosomes before meiotic segregation. Nat Cell Biol 15, 786–796.

Matsuo, Y., Maurer, S.P., Yukawa, M., Zakian, S., Singleton, M.R., Surrey, T., and Toda, T. (2016). An unconventional interaction between Dis1/TOG and Mal3/EB1 in fission yeast promotes the fidelity of chromosome segregation. J Cell Sci 129, 4592–4606.

Maurer, S.P., Bieling, P., Cope, J., Hoenger, A., and Surrey, T. (2011). GTPgammaS microtubules mimic the growing microtubule end structure recognized by end-binding proteins (EBs). Proc Natl Acad Sci U S A 108, 3988–3993.

Maurer, S.P., Cade, N.I., Bohner, G., Gustafsson, N., Boutant, E., and Surrey, T. (2014). EB1 accelerates two conformational transitions important for microtubule maturation and dynamics. Curr Biol 24, 372–384.

Nithianantham S., Cook B., Chang F., Al-Bassam J. (2018) Structural Basis of Tubulin Recruitment and Assembly by Tumor Overexpressed Gene (TOG) domain array Microtubule Polymerases. co-submitted..

Okada, N., Toda, T., Yamamoto, M., and Sato, M. (2014). CDK-dependent phosphorylation of Alp7-Alp14 (TACC-TOG) promotes its nuclear accumulation and spindle microtubule assembly. Mol Biol Cell 25, 1969–1982.

Sato, M., and Toda, T. (2007). Alp7/TACC is a crucial target in Ran-GTPase-dependent spindle formation in fission yeast. Nature 447, 334–337.

Sato, M., Vardy, L., Angel Garcia, M., Koonrugsa, N., and Toda, T. (2004). Interdependency of fission yeast Alp14/TOG and coiled coil protein Alp7 in microtubule localization and bipolar spindle formation. Mol Biol Cell 15, 1609–1622.

Schindelin, J., Arganda-Carreras, I., Frise, E., Kaynig, V., Longair, M., Pietzsch, T., Preibisch, S., Rueden, C., Saalfeld, S., Schmid, B., et al. (2012). Fiji: an open-source platform for biological-image analysis. Nat Methods 9, 676–682.

Telley, I.A., Bieling, P., and Surrey, T. (2011). Reconstitution and quantification of dynamic microtubule end tracking in vitro using TIRF microscopy. Methods Mol Biol 777, 127–145.

Thawani A, Kadzik RS, Petry S. (2018) XMAP215 is a microtubule nucleation factor that functions synergistically with the γ-tubulin ring complex. Nat Cell Biol. 20(5):575–585.

Widlund, P.O., Stear, J.H., Pozniakovsky, A., Zanic, M., Reber, S., Brouhard, G.J., Hyman, A.A., and Howard, J. (2011). XMAP215 polymerase activity is built by combining multiple tubulin-binding TOG domains and a basic lattice-binding region. Proc Natl Acad Sci U S A 108, 2741–2746.

